# From the origin and molecular diversity of the amastins, to the origin and diversity of intracellular parasitism from human Trypanosomatids

**DOI:** 10.1101/2021.11.08.467677

**Authors:** Alejandro Padilla

## Abstract

The large families of amastins from *Leishmania donovani, L. infantum, L. major, L. braziliensis* and *Trypanosoma cruzi* are strongly associated with the evolution of intracellular parasitism of rich cells in human MHC.1 molecules such as the macrophages, dendritic cells, and Langerhans cells by these parasites, recognize the MHC-1 molecules as host receptor. The internalization and transport of the paraste in the cytoplas of infected cell is facilitated by the MHC-1 recycle and endosome formation drag and transport the parasite in the cytoplasm of infected cell. The microbody amastins participate as coreceptor potency the infection, the tropism of *L. major* and *L. braziliensis* by the cells from the skin is facilitated by two molecular interactions, the first molecular interaction is faclitated by the amastins interact the human MHC-1 molecules, and the second molecular interaction is facilitated by the numerous microbody amastins; which also participate in the biogenesis of the small prasitophorous vcuole from *L. major*, and large parasitophorous vacuole from *L. braziliensis*.

All amastins from these parasites developed deactivation domains, in different grade *L. donovani* develop an amastin surface coat specialized in deactivation of infected macrophages heavily glycosylated developed 38 amastins with 38 glycosylation Asp. N-Glycosylation sites and 45 N-glucosamina glycosylation sites, whereas *L. infantum, L. major* and *L. braziliensis* developed one half of glycosylated amastins in asparagine N-glycosylation sites, and *T. cruzi* did not developed none glycosylated amastin.

The amastins surface coat from *L. donovani* is rich in phosphorylation sites, developed 45 amastins with 45 casein kinase II phosphorylations sites, and 48 amastins with 48 protein kinase phosphorylation sites. *L. infantum, L. braziliensis*, and *T. cruzi* developed 32, 42, and 8 amastins, with 94, 114, 21 casein kinase II phosphorylation sites; in similar way developed 35,38, 11 amastins with 89,78, and 22 protein kinase phosphorylation sites. The family of amastins from *L. donovani* develop 137 phosphoserines. and 128 phosphothreonine, *L. major* developed 14 phosphoserine and 4 phosphothreonine; *L. infantum* 1 phophoserine and 7 phosphothreonine; *L. braziliensis* did not developed phosphoserine and phosphothreonine and *T. cruzi* 4 phosphoserine and 4 phosphothreonine. The results show that amastin surface coat is equiped with numerous phosphorylations sites atractive for phosphohrylases from the infected host contribute with the dephosphorylation and deactivation of infectetd host cells.

The amastins from *L. major* develop a membrane amastin with laminin G domain, which can interact with the collagen and heparin sulfate proteoglycan sites from the extracellular matrix of the skin tissue. Furthermore develop 14 amastins with tyrosine sulfation site, evade the activation of receptor of chemokines and the activation of the immune response by chemokines.

There is an alternative mechanism of polarization of the immune response from protective TH1 to non protective TH2.

The parasite nutrition is mediated by amastins that dissimilate the MHC-1 molecules and other subsets of proteins, the dissimilation products can be translocated through of the parasite cell membrane and employed as nutrient source.

## Introduction

Leisshmaniasis is a neglected human parasitic disease that affects the tropical and subtropical regions of the world [1]. These parasitosis affects 12 millions people, with 2 millions of new cases appearing every year, and threat approximately 350 millions people in the endemic area [1]. The leishmaniasis is a spetrum of disease caused by the parasite protozoan *Leishmania* that course from simple cutaneous ulcers to the mortal visceral leishmaniasis [2]. Currently, there are no vaccines to prevent this conditions [3]; and reliable drugs that can be used for the successful treatment of the infection [4].

The Chagas disease or trypanosomiasis Americana is caused by parasite protozoan Trypansosoma cruzi, is one of the most important parasitosis in Latinoamerica affect from 12 to 20 million of people, and 50000 people dead yearly by complications associated with Chagas disease, cause cardiomegaly, megacolon, and megaesophogus [5].

The amastins were first identified in *T. cruzi*, and were called amastins due to their upregulated expression associated with the differentiation of the amastigote stage [6]. The amastins are positioned in different chromosomes, beside the genes encode tuzins, which were also identified in *T. cruzi*, and described as rare proteins of unknown fucntion and uncertain cellular location [7]. The amastins from *Leishmania infantum, L. major*, and *L. mexicana*, are also known and are associted with the differentiation of the amastigote stage in the parasite life cycle [85], and are stritcly over expressed by acidid pH, and regulated by DNA elements located in the 3’untranslated region of their mRNA [8-9]. The promising immunologial properties of amastins came to light when amastins from *L. major* became the best antigen to induce immunity against the experiemental infection with *L. major* among the 100 novel antigens tested in a mice model [10]. The amastins from *L. major* also show the protective capacity employ DNA vacccine, using amastins fused to HSV-1 VP22 and EGFP in BALB/c mice model [11]. The amastins from *L. infantum* incoroporated in different adyuvant systems induce protection against visceral leishmaniasis [12]. The immunogenecity and protective immunity also was show with multi-epitopes DNA prime-protein, boosst vacccines encoding amastins -kmp-11, kmp11-Gp63, and amastin-Gp63 protect against visceral leishmaniasis [13].

The relevance of amastins in the amastigote was show by amastin knock down in *L. braziliensis*, affected the interaction of the parasite with the macrophage and the viability of the amastigote [14].

In this work, the families of amastins from human parasites *Trypansoma brucei, T. brucei gambiense*, and the animal parasites *T brucei strain 427, T. vivax, and T. congolense;* were investigated and compared with the families of amastins from the human para<u>sites</u> *L. donovani, L. infantum, L. major, L. braziliensis, and T. cruzi*, employing the bioinformatical tools; in order to know the genomic response, that developed the evolutionary strategies employed for the origin and construction of the molecular diversity of the repertoire of amastins from these parasites; deduce the role played by the amastin domains in the infection of host cells, internalization and transport of the parasite in the infected cell; tropism of the parasites by the cells from the skin or internal organs; biogenesis of the parasitophorous vacuole (PV); evasion of the immune response; and parasite nutrition; finally identify the molecular and functional diversity of the families of amastins that developed the intracellular parasitism from *L. donovani, L. infantum, L. major, L. braziliensis*, and *T. cruzi*; and their role in the pathogensisis of their respective human parasitic disease.

## Material and Methods

### Amino acid sequences of amastins

The families of amastins investigated from the human extracellular parasite *T. brucei* [15], and *T. brucei gambiense* were acquiered from their respective protein data base. As well as the amastins from the animal Trypansomatids *T. brucei strain 427*; *T. vivax* and *T. congolense*, were obtained from their respective protein database. The amastins from *L. major* [16], *L. infantum* [17], *L. donovani* [EBI site], *L. braziliensis* [17] and *T. cruzi* [18], were acquired from repective database, copy and pasted in word processor, for analyses with the bioinformatical tools.

### Bioinformatical tools

The mature amastins from these parasites were analyzed utilizing nine bioinformatical tools from expasy site (https://www.expasy.org/). The amino acid sequence alignment was performed in the European Bioinformatical Institute EMBL-EBI site by clustal W program (http://www.ebi.ac.uk/clustalw/) [19-21]. Hydrophobic patterns in scale were analyzed utilizing the ProtScale program (http://ca.expasy.org/tools/protscale.html) [21].

The number of transmembrane segments that compose the amastins were established using “DAS” transmembrane prediction program (http://www.sbc.su.se/~miklos/DAS/)

[22] Isoelectric point (pI) and molecular weight (MW) of amastins were assessed by the compute pI/MW tool (http://ca.expasy.org/tools/pi_tool.html) [21, 23].

The amino acid repeats were investigated in REP-site (http://www.embl-heidelberg.de/~andrade/papers/rep/search.html) [24]. The similarity of the amastins with the amino acid sequence reported in the gene bank was investigated by blastp analysis (http://www.ncbi.nlm.nih.gov/BLAST/) [25]. The cellular targets of the amastins were investigated by analyzing the N-terminal region, and identifying the domain of signal anchor, signal peptide, or non seretory proteins (http://www.cbs.dtu.dk/services/LipoP/) [26-27]. The prosite site was used to identify the domains, signatures, and fucntionl sites of the amastins (http://ca.expasy.org/prosite/) [21]. The human MHC-1 motifs from amastins were identified by a program developed by [22].

### Parasite genome project

In this work was analyzed the amastins from *T. brucei* genome (Berriman et al 2005), [http://www.genedb.org/genedb/tryp/index.jsp]; as well as the amastins from the human trypansosomatid *T. brucei gambiense* [http://www.genedb.org/genedb/tgambiense/]. As well as the amastins were searched in the animal trypansomatid *T. brucei strain 427* [http://www.genedb.org/genedb/tbrucei 427]; *T. vivax* [http://www.genedb.org/genedb/tvivax/]; *and T. congolense* [http://www.genedb.org/genedb/tcongolense/].

Sequences of the amastins from *L. donovani* genome (EBI site) (https://www.ebi.ac.uk/ebisearch/search.ebi?db=proteinSequences&query=L.%20donovani%20genome%20amastin&size=15&requestFrom=searchBox&page=4)

Sequences of the amastins from the chromosome 36, 34, 31, 30, 28, 24, and 8 of *L*.*major* were analyzed as a part of the *Leishmania* genome network (LGN), (http://www.sanger.ac.uk./project/L_major/), as well as the amastins of the chromosomes 31, 30, 28, 24, 20,18,13, and 8 from *L. braziliensis* genome [http://www.genedb.org/genedb/lbraziliensis/]. The amastins of the chromosome 34, 31, 30, 29, 24, and 8 from *L. infantum* genome [http://www.genedb.org/genedb/lbraziliensis/]. The amastins of *L. donovani* were acquired from EBI site [https://www.ebi.ac.uk/ebisearch/search.ebi?db=proteinSequences&query=L.%20donovani%20genome%20amastin&size=15&requestFrom=searchBox&page=4].

The amastins from *T. cruzi* analyzed here were obtained from the *T. cruzi* genome project [http://www.tigr.org/tdb/e2k1/tca1/]. All the trypanosomatid genome projects analyzed in this work were economically supported by the Wellcome Trust Sanger Institute.

## Results and Discussion

### The families of amastins

The search of amastins in the human extracellular *T. brucei* genome show that this parasite developed a couple of genes enconde for intracellular amastins, compose by four transmembrane segments, one acidic and one basic (Tab. 1).

**Tab. 1.** Deduced role of the amastins versus *L. donovani, L. infantum, L. major, L. braziliensis, T. cruzi, T. brucei*. Show that *L. major* perform the most complex genomic response, followed by *L. braziliensis, L*.*donovani, L. infantum* and *T, cruzi*. The amastins from *T. brucei* did not perform genomic response, The amastins from all these parasites perform infection of host cells, internalization and transport of the parasite in the cytoplasm of infected cell, polarization of the immune response from protective TH1 to non protective TH2, and parasite nutrition mechanism. The amastins from *T. cruzi* perform the less complex genomic response, achieve additional acid resistant mechanism by basic amastins. Followed by the genomic response from *L*.*infantum* that achieve additional coreceptor for infection single microbody amastin, and molecular mimics by amastins ALDH, The genomic response by *L. donovani* develop additional co receptor for infection, acid resistant mechanism by basic amastins, *L. braziliensis* achieve additional co receptor for infection with 3 microbody amastins, tropism of the parasite by the cells from the skin, and biogenesis of the large and multiple parasitophorous vacuole. Finally the *L. major* perform additional co receptor for infection with 9 microbody amastins, tropism of the parasite by the cells from the skin, and biogenesis of small and individual PV.

The search of amastins in other African trypanosomatids as the human extracellular *T. brucei gambiense* show that developed single amastin gene. The animal extracellular *T. brucei strain 427, T. vivax*, and *T. congolense* didn’t developed amastins genes in their genomes. By the contrary the results show that large families of amastins were developed by human intracellular parasites *Leishmania* donovani develop a family compose by 50 mature amastins (Fig. 1), *L. infantum* developed a family compose of 40 mature amastins (Fig. 2), *L: major* develop a family compose by 54 mature amastins (Fig. 3); *L. braziliensis* developed a family compose by 47 mature amastins (Fig. 4); and *T. cruzi* develope a family compose by 12 mature amastins (Fig. 5). The results show the parasites that developed few amastins as *T. brucei*, and *T. brucei gambiense* and the animal African trypanosomatids did not developed the intracellular parasitism, developed the extracellular parasitism. The large families of amastins such as *L. donovani, L. infantum, L. major, L. braziliensis* and *T. cruzi* are associated with the differentiation of the amastigote stage specialized in develop the intracellular parasitism of human cells from these parasites (Tab. 1).

**Fig. 1.**
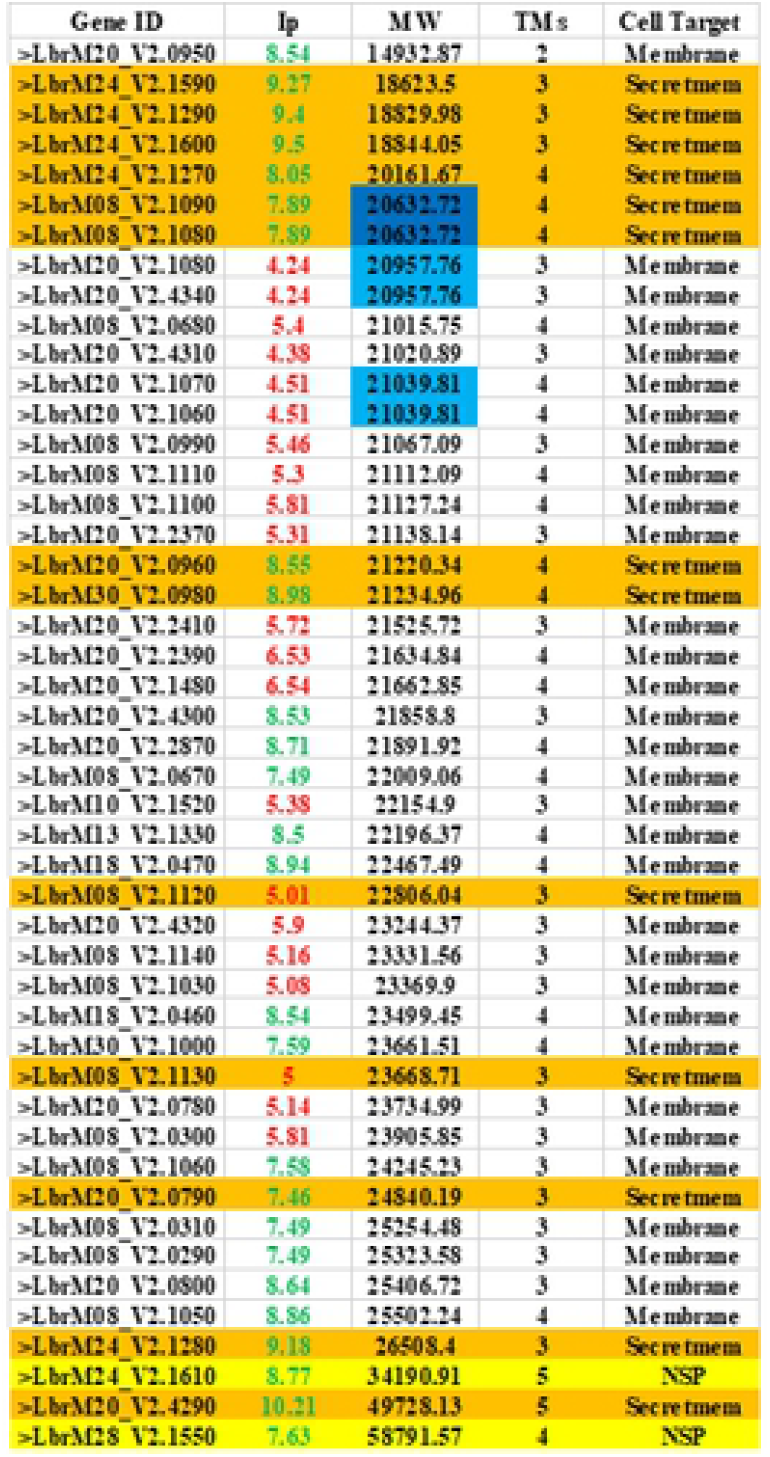
Diversity of the family of amastins from *L. donovani* compose by 50 mature proteins: Column A= ID number of the amastins. Column B=Isoelectrical point. Column C=Molecular weight. Column D=Transmembrane segments. Column E= Cellular target. The amastins are ordered by molecular weigh (MW), show that diversity of these molecules vary from 15.36 KD to 22.73 KD; 12 of the most large amastins are non secretory proteins (NSP) are 13 NSP with varyable MW= 23.25 KD to 57.85 KD (yellow color), contain the 3 atypical amastins of eight transmbenrane segments. 14 amastins developed the domain of signal peptide, pass by the secretory pathway and are targeted in the parasite cell membrane (orange color). 23 amastins developed the domain of signal anchor, and are targeted in the parasite cell membrane (white color). 20 amastins present acidic iP (red numbers), and 30 are basic amastins (green numbers), developed acid resistant mechanism. The diversity can be see in the number of transmembrane segments (TMs.) 43 amastins developed 4 TMs., 3 amastins developed 3 TMs., 3 amastins developed 8 TMs., single amastins did not developed TMs.

**Fig. 2.**
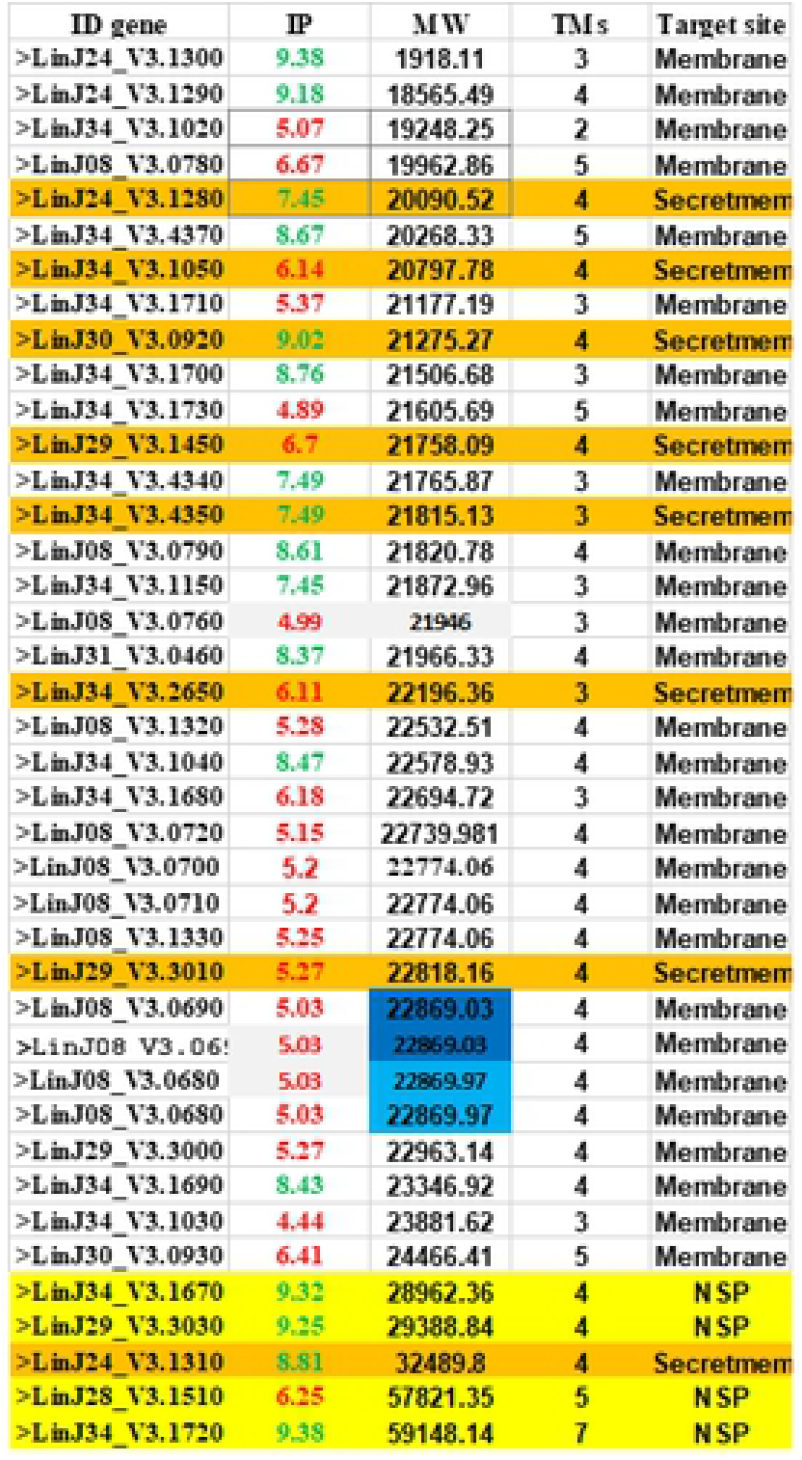
Diversity of the family of amastins from *L. infantum* compose by 40 mature proteins: Column A= ID number of the amastins. Column B=Isoelectrical point. Column C=Molecular weight. Column D=Transmembrane segments. Column E= Cellular target. The amastins are ordered by MW, show that diversity of amastins vary from 19.18 KD to 24.98 KD; and the exception 5 large amastins that vary from 28.96 KD to 57.14 KD which are mainly NSP (yellow color). There are 8 secreted amastins that developed the signal peptide domain and enter to the secretory pathway and the mature proteins are targted in the parasite cell membrane (orange color). There are 28 membrane amastins (white color); thereare two couples of identical amastinas (blue color). The isolelectrical point show that there are 17 basic amastinas (green color), and 23 acididc amastins (red color). There are 23 amastins composed by 4 transmembrane segments (TMs), followed by 10 amastins of 3 TMs.; 5 amastins of 5 TMs. Single amastins developed 7 TMs. and 2 TMs.

**Fig. 3.**
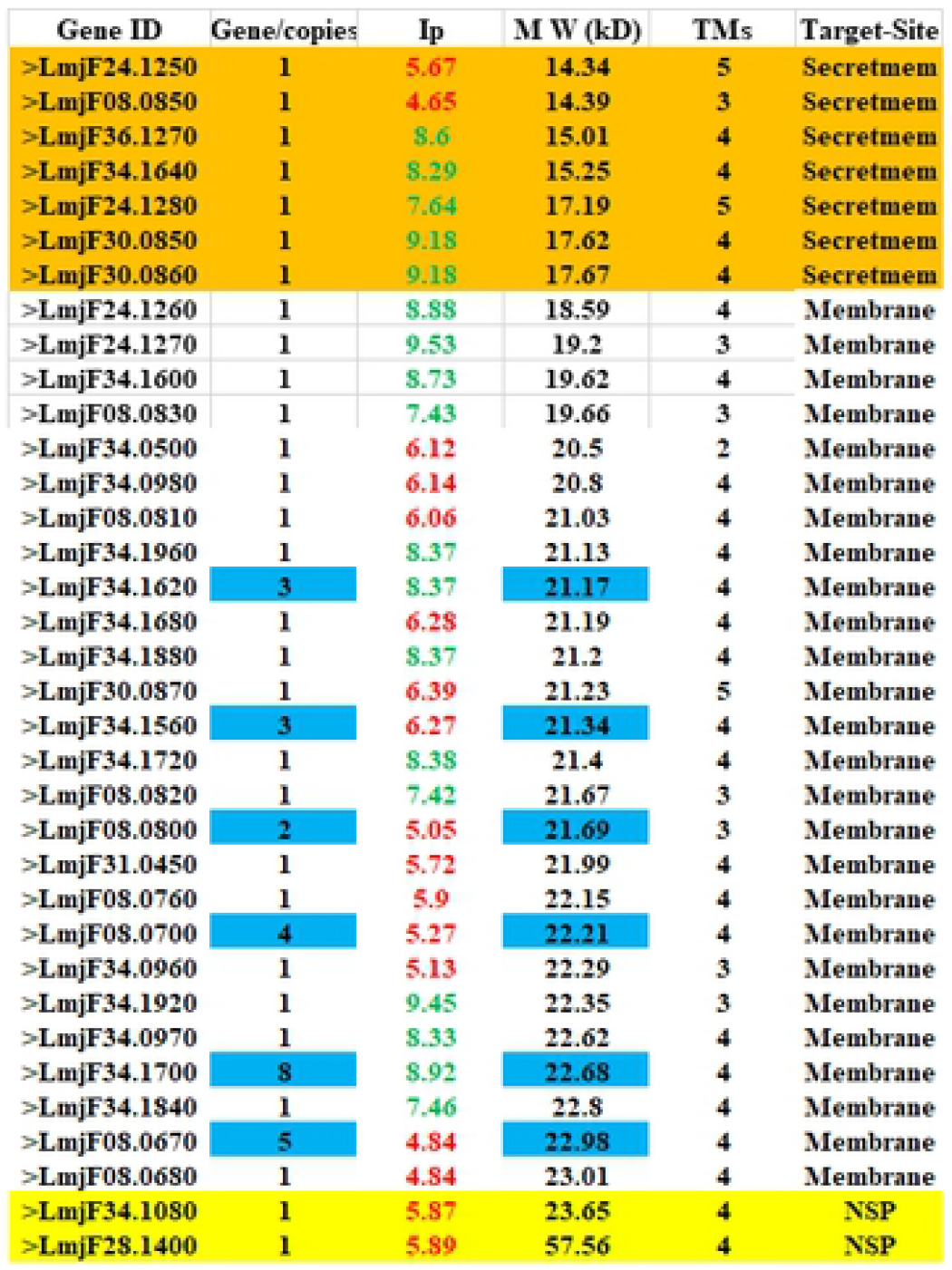
Diversity of the family of amastins from *L. major* compose by 54 mature proteins: Column A=Gene ID number. Column B=# Gene copies C=Isoelectrical point. Column D=Molecular weight. Column E=Transmembrane segments. Column F= cellular target. The amastins are ordered by MW show that vary from 14.34 KD to 23.66 KD, with the exception of single large amastin of 57.56 KD. There are 6 amastins in multicopy gene (Blue color). The isoelectrical point show 27 acidic amastins (red color). and 27 basic amastins (green color). There are 2 non secretory proteins (yellow color), there are 7 secretory amastins targeted in the parasite cell membrane (orange color): and there are 45 membrane amastins (white color). The diversity can be appreciated in the number of TMs, there are 42 amastins with 4 TMs; 8 amastins with 3 TMs; 3 amastins with 5 TMs; and single amastin with 2 TMs.

**Fig. 4.**
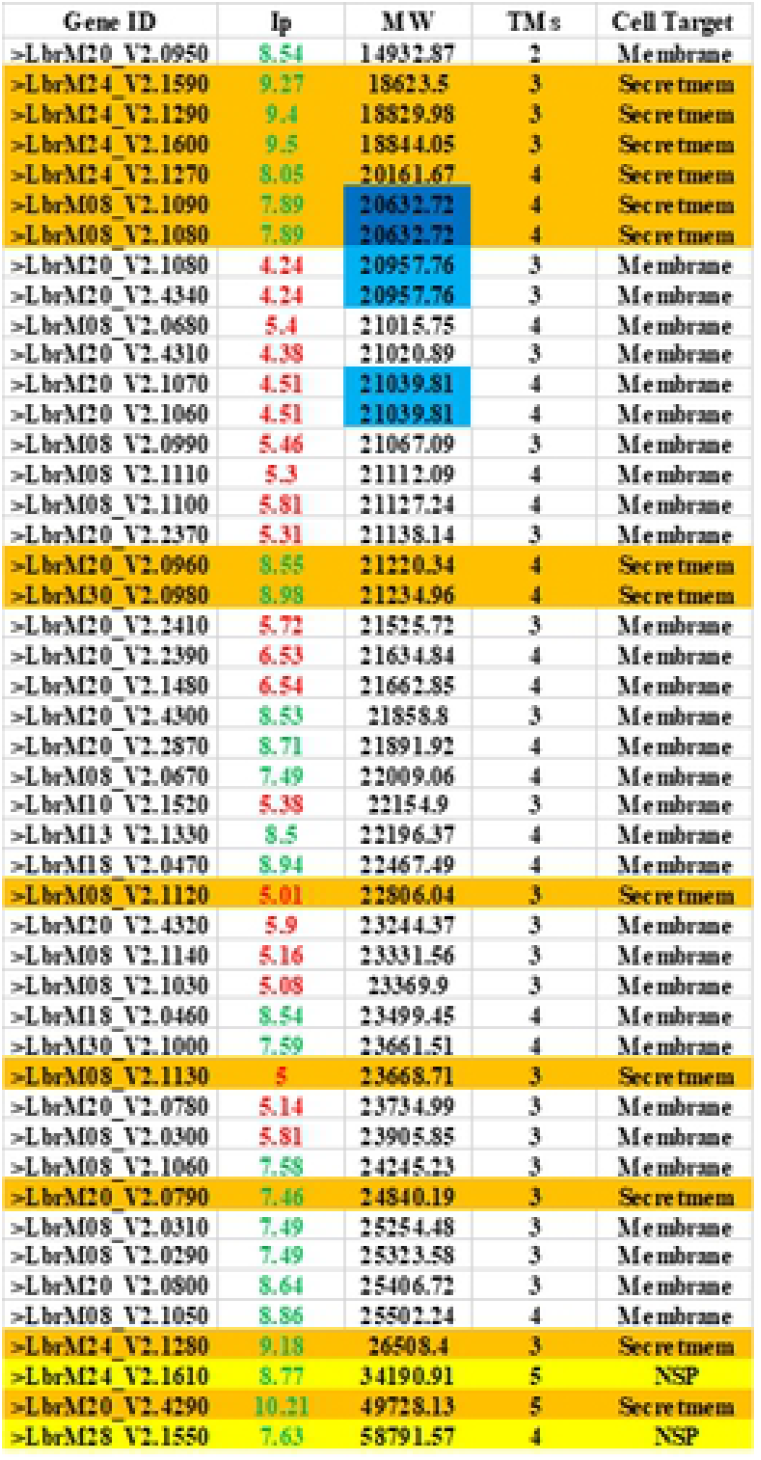
Diversity of the family of amastins from *L. braziliensis* compose by 47 mature proteins; Column A= ID number of the amastins. Column B=Isoelectrical point. Column C=Molecular weight. Column D=Transmembrane segments. Column E= cellular target. The amastins are ordered by MW show that amastins vary from 14.93 KD to 26.50 KD with exeption of three large amastins that vary from 34.19 KD to 58.79 KD. There are three couples of identical amastins (Blue color). The isoelectrical point show that developed 26 basic amastins (green color) and 21 acidic amastins (red color), There are two NSP or intracellular amastin (yellow color), there are 13 secreted amastins that developed the signal peptide and pass by the secretory pathway and the mature protein is targeted in the parasite cell membrane (orange color), there are 32 membrane amastins that developed the domian of signal anchor in the N-terminal (white color). The polymorphism also can be appreciated in the number of TMs., show that 24 amastins developed 3 TMs., 20 amastins developed 4 TMs., 2 amastins developed 5 TMs., and single amastin develop two transmembrane segments.

**Fig. 5.**
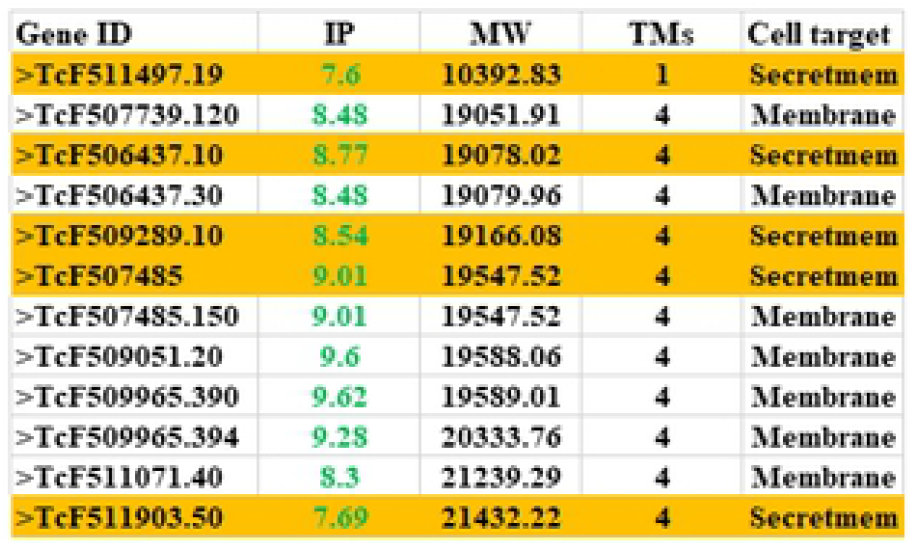
Diversity of the family of amastins from *Trypanosoma cruzi* compose by 12 mature amastins: Column A= ID number of the amastins. Column B=Isoelectrical point. Column C=Molecular weight. Column D=Transmembrane segments. Column E= Cellular target. Ordered by molecular weight show that amastins vary 19.05 KD to 21.43 KD, with exception of single small amastin of 10.39 KD. The isoelectrical point show that all amastins are basic develop acid resistant mechanism (green color). *T. cruzi* just developed 5 secretory amastins that enter to the secretory pathway and the mature amastins are targeted in the parasite cell membrane (orange color), and 7 membrane amastins (white color). There are 11 amastins that developed 4 TMs., and single small amastin developed single TMs.

### Origin of the molecular diversity of amastins

The analyses of the repeats of the amino acid sequences of the amastins, show that amastins genes where shaped by recombination of two small fragments, shape one halve, which by gene replication and recombination of two halves shape mature gene of 198 amino acids (Fig. 6a). Or can be shaped by recombination of four small fragments of DNA encode for around of 21 amino acids shape mature gene of around of 198 amino acids, compose by four transmembrnane segments (Fig. 6b). The enlargement of the repertoires of amastins from *L. major, L. infantum, L. donovani, L. braziliensis* and *T. cruzi*, were made by gene copy and independent maturation of the amastin genes, shape the diversity of the large repertoires of amastins (Figs. 1-5).

**Fig. 6.**
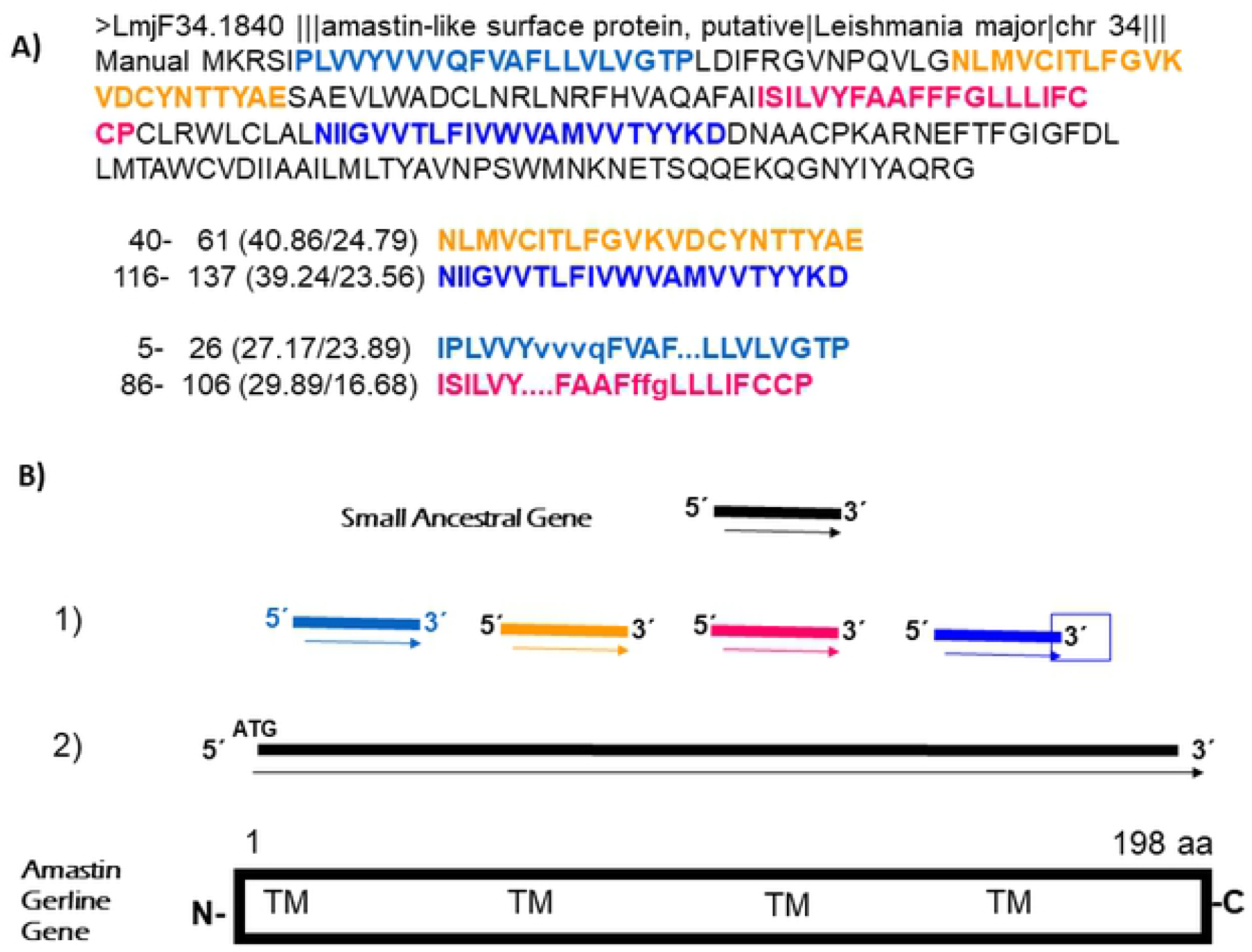
**a)** Amino acid sequence repeats, show that amastin >Lmjf34.1840 was shaped by four fragments of DNA of around of 21 amino acids, **b)** The recombination of four fragments of DNA shape the mature gene encode for amastins of 198 amino acids (1, 2),

### Analyses of transmembrane segments and variable regions of amastins

The amastins are hydrophobic molecules (Fig. 7a), generally compose by four transmembrane segments (Fig. 7b), shows the analyses of amastin (isoform LmjF08.0750). As observed the firsts transmembrane segments of the amastins starts from the amino acids 7-10, and ends with the amino acids 24-29. The second transmembrane segment starts from the amino acids 80-87 and ends with the amino acids 102-106. The third transmembrane segments starts from the amino acids 108-112 and ends with the amino acids 125-131. The fourth transmembrane segments starts from the amino acids 147-184 and ends with the amino acids 171-198 (Fig. 7 b).

**Fig. 7.**
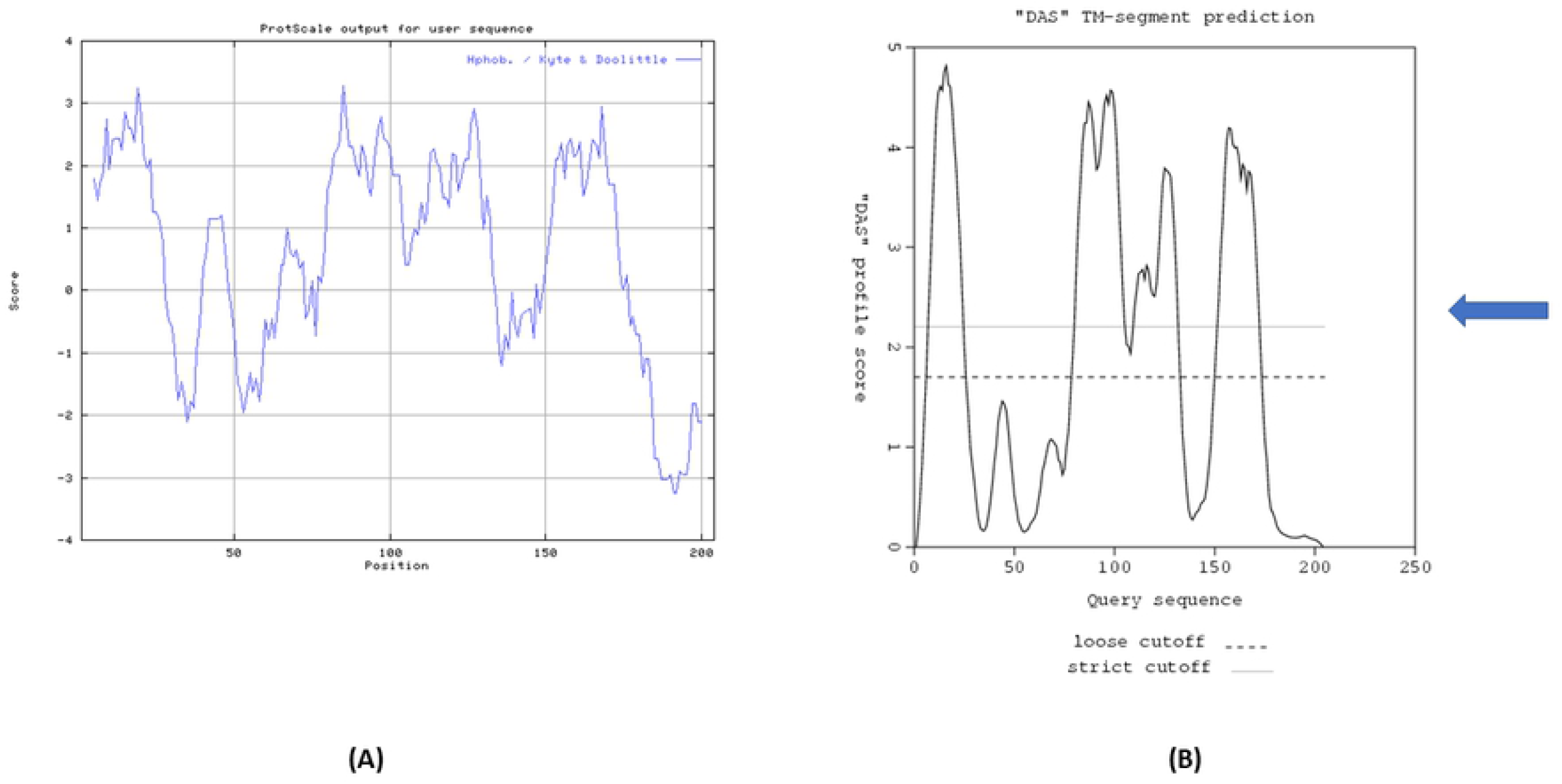
**a)** Hydrophobic patterns, show that amastins are hydrophobic molecules compose by four hydrophobic regions, and three or four hydrophilic regions, **b)** Analyses of transmembrane segments show that amastins is compose mainly by four transmembrane segments (Arrow indicate the cut off for transmembrane segments = 2.2).

The results furthermore show that amastins develop 3 variable regions of around 50 amino acids positioned in the hydrophilic regions (Fig. 7a): first variable region (amino acids 30-80) after the first transmembrane segment in the N-terminal (Fig. 7 a), second variable region of around 30 amino acids (amino acids 125-155) between the third and fourth transmembrane segments (Fig. 7a), and the third variable region positioned in the C-terminal of these molecules (Fig. 7a). The first variable region tends to create diverse active sites such as sulfation sites, cAMP- and cGMP-dependent protein kinase phosphorylation sites, protein kinase C phosphorylation sites, and tyrosine kinase phosphorylations sites. Further, it also creates diverse MHC-1 motifs by the human MHC-1 molecules (Fig 7a). The second variable region develops from 1 to 4 N-asparagine glycosylation sites, creates diverse MHC-1 motifs by the human MHC-1 molecules (Fig. 7a). The third variable region creates diverse domains of the C-terminal from these molecules, as multiple MHC-1 motifs and the microbody domain (Fig. 7 a).

The amastins from *L. infantum, L. braziliensis*, and *T. cruzi* show the four transmembrane segments with four variable regions (Data not shown). The repertoire of amastins from *L. braziliensis* possesses 25 amastins with three transmembrane segments, 21 amastins with four transmembrane segments, two amastins developed five transmebrane segments, and single amastin developed two transmembrane segments. The cause of there are more amastins with three transmembrane segments, that amastins with four transmembrane segments in *L. braziliensis*, is the fusion of hydrophobic amino acids of the second and third transmembrane segments, form membrane amastins of three transmembrane segments (Fig. 4).

### Analysis of the N-terminnal domains of the amastins

The analyses of the N-terminal of the amastins from these parasites reveal polymophism of the domains, posses the signal anchore of membrane amastins, signal peptide of secreted amastins, and hydrophilic region of non secretory proteins (can be intracellular amastins) (Fig. 8 a-c).

**Fig, 8.**
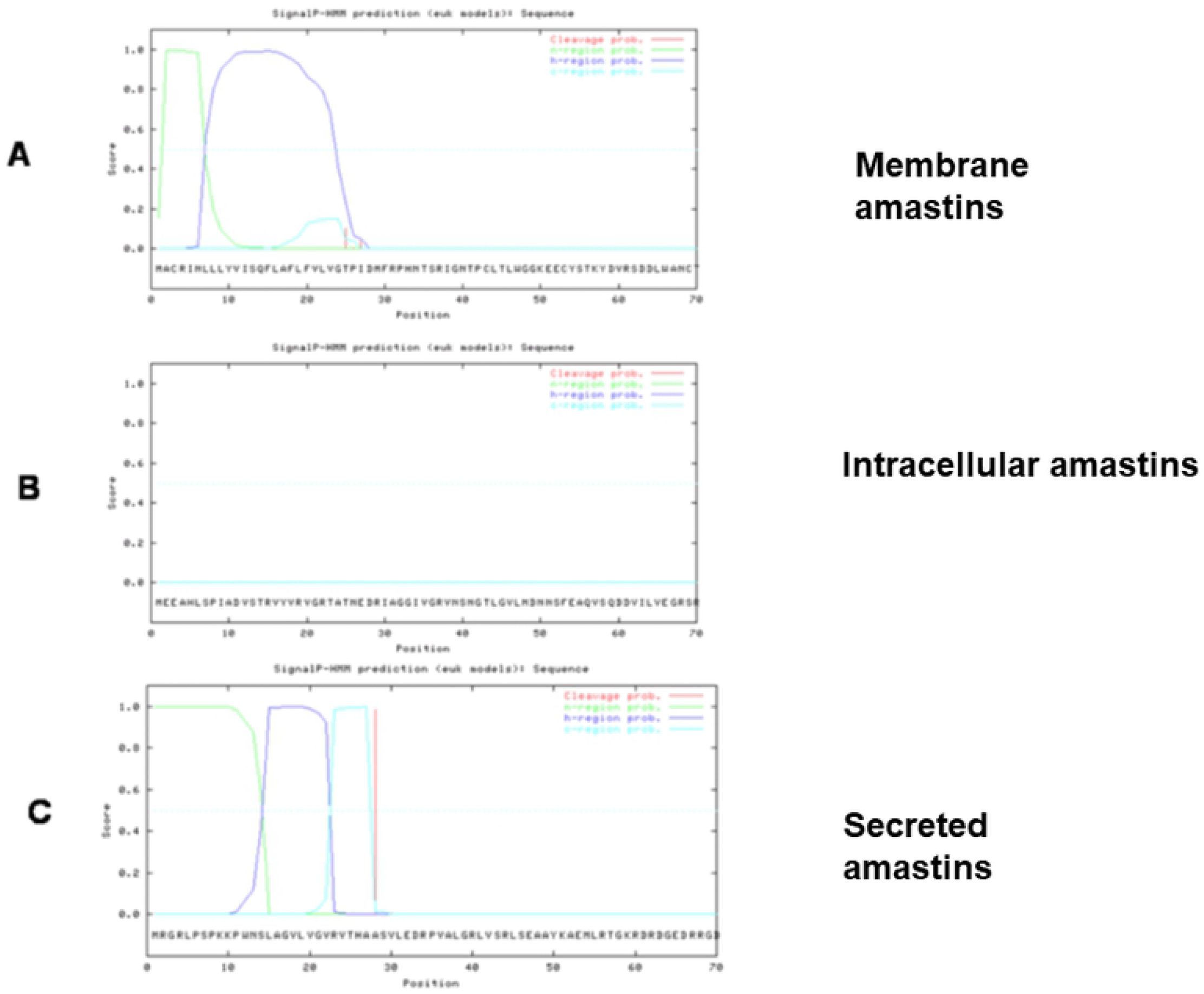
Analyses of N-terminal region of the amastins show polymorphism. **a)** Membrane amastins developed the domain of signal anchor in the N-terminal, **b)** Non secretory proteins or intracellular amastins develop hydrophilic N-terminal. **c)** Secretory amastins develop the domain of signal peptide in the N-terminal, show the region of dissimilation of amino acids of the signal peptide, shaped by neutral amino acids.

Signal anchore domain were identified in the N-terminal region of the amastins. Our findings indicate that these molecules enter to the secretory pathway are localized in the cell membrane of the amastigote stage. They can possess signal peptide in the N-terminal, which is indicative that the polypeptide pass by the secretory pathway; as the secreted amastins are hydrophobic molecules, remain attached to the parasite cell membrane by three to four transmembrane segments. The absence of the signal anchore or signal peptide of the N-terminal, and development of hydrophilic sequence in the N-terminal is characteristic of the non secretory proteins (which can be intracellular amastins) (Fig. 8 a-c).

### Analysis of membrane amastins

Majority of amastins are localized in the cell membrane of these parasites, conserving the hydrophobic regions signal anchor domain. This domain is recognized in the nascent polypeptides by the signal recognition particle (SRP), which translocate the polypeptide to the lumen of the endoplasmic reticulum (ER) to follow their maturation process in the secretory pathway [23], and to finally be localized in the parasite surface membrane (Fig. 8a).

### Analysis of non secretory amastins (intracellular amastins)

Results show that the intracellular amastins from these parasites, were generated by the addition of different fragment of DNA, encoding the large hydrophilic fragment of around 350 amino acids which is positioned in the N-terminal region ancestor gene of the cell surface membrane amastins, and shape the new genes that encode for intracellular amastins. The hydrophilic region prevensts the binding of the nascents polypeptide to SRP. The hydrophilic region prevents the biding of the nascent polypeptide to SRP and the access of the new polypeptide to the secretory pathway, and hence, the peptide is redirected to the additional intracellular compartment of the parasite cell (Fig. 8b).

### Analyses of secretory amastins

Our analyses revealed the evolutionary strategy employed to create secretory amastins from these parasites, was the generation of the signal peptide by random mutations in the signal anchor region of the membrane amastin genes; changing the amino acid sequence of the protein that codes for the cleavge site to remove the signal peptide from the secreted amastins. The position of the cleavage sites from the secreted amastins vary from the amino acid 24 to 29, conserve the rule-3, -1 of the cleavage site which should exist an small and neutral region of amino acids employed for the cleavage of the signal pepide by the signal peptidase. The amino acid occupying the -1 position of the signal peptide from secreted amastins are alanine, glycine, and serine; and in the position -3 are the amino acids alanine, glycine, and valine. The variable sequence and positions -3 are the amino acid alanine, glycine, and valine. The variable sequence and positions of the cleavage sites of the signal peptides indicate that these molecules were shaped by different and independent evolutionary phases of the creation of the repertoire of amastins (Fig. 8 c).

### The family of amastins from *Leishmania donovani*

The results show that family of amastins from *L. donovani*, is compose by 50 mature amastins (Tab. 1), the amastins are ordered by isoelectric point, show that 20 amastins are acididc with variable Isoelectrical point (Ip) from 4.37 to Ip 6.79, and 30 amastins are basic vary Ip from 7.45 to Ip 11.37. The molecular weight (MW) of the amastins show that diversity, vary from 15.36 KD to 57.85 KD (Fig. 1).

The analyses of the transmembrane segments that compose the amastins from *L. donovani*, show that 44 amastins are compose by 4 transmembrane segments, 2 amastins are compose by three transmembrane segments, three atypical and large amastins compose by 8 transmembrane segments, and one hydrophilic segment without transmembrane segments, which could be employed to shape large intracelllular amastins of four transmembrane segments (Fig. 1)

Amazingly 10 of 13 intracelular amastins show large molecular weight vary from 24.46 KD to 57.8 KD. 10 of these molecules show basic iP. The three amastins of 8 transmembrane segments did not developed none special domain. These atypical molecules can participate regulate the differentiation of the promatigote stage to amastigote stage by acidic pH (Fig. 1).

### Amidation sites of the amastins from *L. donovani*

The analyses of the amiadation sites of the amastins from *L. donovani* show that this parasite develop 3 amastins with 4 amidation sites. The biological relevance of the amidation sites show that play important role in the translocation of the parasite from the internal leaf to the nucleus of infected cell; employing the molecular machinary of G proteins coupled receptor, transport the parasite from the cell membrane through of the cytoplasm to the nucleus of the infected cell [92]. However the results show that developed a four amidation sites thereby this mechanism did not play important role in the transport of the amastigote through of the cytoplasm of infected cell, which should be occur by the MHC-1 recycle and endosome formation, drag and transport the parasite in the cytoplasm of infected cell (Tab. 2).

**Tab. 2.** Domains and active sites of amastins versus *L. donovani, L. infantum, L. major, L. braziliensis, T. cruzi*. Amidation sites *L. infantum* developed most amiadation sites 7 with 7 active sites, *T. cruzi* did not develop none amidation site. ASN-Glycosylation N-glycosylation site *L. donovani* develop 38 amastins with 38 active sites, *L. infantum, L. major* and *L. braziliensis* developed respectively 21, 23 and 25 amastins with 31, 58, and 47 active sites. *T. cruzi* did not developed ASN-Glycosylation N-glycosylation site. N-linked (GlcNAc…) asparagine, *L. donovani* develop 45 amastins with 45 active sites, the rest of parasites did not developed none N-linked (GlcNAc…) asparagine reveal that amastins from *L. donovani* are the most N-glycosylated amastins. cAMP and cGMP dependt protein kinase phosphorylation site *L. braziliensis* develop 13 amastins with 13 active sites and *T. cruzi* develop 3 amastins with 3 active sites. Casein kinase II phosphorylation site *L. braziliensis* develop 40 amastins with 114 active sites and *L. major* the minor develop 16 amastins with 28 active sites. Protein kinase C phosphorylation site *L. donovani* develop 48 amastins with 48 active sites; *T. cruzi* develop 11 amastins with 22 active sites, and *L. major* develop 13 amastins with 27 active sites. Tyrosine kinase phosphorylation site 1, just *L. donovani* develop 2 amastins with 2 active sites. Phosphoserine and Phosphotreonine *L*.*donovani* develop respectively 137 and 128 *L. braziliensis* develop 0. Microbodies C-terminal targeting signal *L. major* develop 9 amastins with microbody domain, *L. braziliensis* develop 3 amastins with 3 microbody domain, and *L. infantum* and *L. donovani* developed single amastins with single microbody domain. N-myristoylation site all parasites developed numerous amastins with numerous active sites. Cell attachment sequence *L. donovani, L. infantum* and *L. major* developed amastins with cell attachment sequences. Aldehyde dehydrogenases cysteine active site just *L. infantum* develope two amastins with this domain. Laminin G domain just *L. major* develop single amastin with this domain. Tyrosine sulfation site just *L. major* developed 14 amastins with this domain.

### Asparagine N-linked glucosamine, and Asparagine-glycosylation site of the amastins from *L. donovani*

Show that developed various monosaccharides, oligosaccharides and their derivative form bonds with different amino acid residues within a protein as result of glycosylation. There are five classes of glycosylation: N-linked, O-linked, C-linked, Phospho glycosylation and glypiation. Every kind of glycosylation imparts a special characteristic to the modified protein as required by its role in cellular process. N-linked glycosylation is common amongst all types as it holds 90% share in total glycosylations. The exposed asparagine residues of a protein are found to form N linked bond with sugars. Any asparagine (N) residue appearing within a consensus pattern of sequence will form N-linked bond with sugars. This modification is processed in endoplasmic reticulum (ER) lumen before exporting the modified protein to the cytoplasm or outside of the cell. In ER lumen dolichol molecule plays a pivotal role in this process. The membrane-bound dolichol molecule has a long chain isoprene whose one end is attached with isoprenoid group and other with saturated alcohol. [87].

The results of this work show that 45 amastins from *L. donovani*, developed asparagine N-linked glucosamine, and 38 amastins developed ASN-glycosylation site. Actually is well know that N-glycosylation from parasistes, virus, bacteria, play important role in the evasion of the innate immune response [25], the antibody response [89], antigenic variation [90], immune response evasion [28] from the human infected host. Thereby the N-glycosylated amastins from *L. donovani* can play important role in evasion of the innate and specific immune response, and antigen variation, allow the installation of the amastigote and the develop of the visceral leishmaniasis (Tab. 2).

### cAMP and cGMP-dependent protein kinase phosphorylation site of the amastins from *L. donovani*

The family of amastins from *L. donovani* developed 10 amastins with 12 cAMP and cGMP-dependent protein kinase phosphorylation active sites. These kinases use serine and threonine as phosphorylation substrate. Reveal that cAMP and cGMP-dependent protein kinase phosphorylation site, can be atractive for the cAMP and cGMP-dependent protein kinases from the infected macrophages, interfere the regular phosphorylation signaling from infected host cell, and contribute with the deactivation of the infected macrophages (Tab. 2).

### Casein kinase II phosphorylation site of the amastins from *L. donovani*

The analyses show that repertoire of amastins from *L. donovani*, develop 45 amastins with 45 casein kinases II phosphorylation active sites. The casein kinase II is a phosphoserine, phosphothreonine phosphorylation protein; but their activity is independent of cyclic nucleotides and calcium. Reveal that casein kinase II phosphorylation site can be atractive for the casein kinase II phosphorylases from the infected host cell, interfere the regular cell signalling and contribute with deactivation of infected macrophages, allow the growth of the amastigotes, and the develop of the visceral leishmaniasis (Tab. 2).

### Protein kinase C phosphorylation site of the amastins from *L. donovani*

The analyses show that amastins from *L. donovani* developed 48 amastins with protein kinase C phophorylation site, furthermore developed numerous phosphoserine (137) and phosphothreonine (128), reveal that amastins from *L. donovani* are widely phosphorylated molecules. These phosphorylation sites can play important role in the hosts parasite relationship, since can be atractive for the protein kinase from the infected macrophage, interfere the regular phosphorylation cascade of the macrophages contribute with the deactivation of the infected cell (Tab. 2).

### Tyrosine kinase phosphorylation site 1 of the amastins from *L. donovani*

The analyses show that repertoire of amastins from *L. donovani* developed 2 amastins with tyrosine kinase phosphorylation sites. Substrates of tyrosine protein kinases are generally characterized by a lysine or an arginine seven residues to the N-terminal side of the phosphorylated tyrosine. Reveal that this Tyrosine kinase can participate in the deactivation of the infected cells being attractive for the tyrosine kinase from the infected macrophage, contribute with the deactivation of the infected cell. (Tab. 2).

### Cell Attachment sequences of the amastins from *L donovani*

The search of cell attachment sequences of the repertoire of amastins from *L. donovani* show, that developed 4 amastins with the minimal sequence of recognition RGD, and 5 amastins with the minimal recognition sequence with the amino acids LDV (Tab. 2).

The cell attachment domain with the minimal recognition sequence RGD can interact the integrins from the extracellular matrix, contribute with the attachment of the parasite to the extracellular matrix, from the infected spleen, liver and bone marrow of the visceral leishmaniasis. The 5 amastins with the minimal recognition LDV typical from integrins from leukocytes, can participate in the infection of the macrophages, attach the parasite to the LDV integrins from the cell membrane of the host cell. Or evasion of the immune response interfere the correct binding of the chemokines with its receptor [29]. Interfere the correct synapsis of activation of the macrophages and T lymphocytes, contribute with the deactivation of the immune cells (Tab. 2).

### Microbody C-terminal targeting signal of the amastin from *L*.*donovani*

The analyses show that amastins from *L. donovani* developed single microbody amastin, similar to *L. infantum*. The microbody amastin can participate in the attachment of the amastigote to the cell membrane from the host cells. The single microbody amastins can not support the biogenesis of the parasitophorous vacuole (PV) [30], nor the tropism of the parasite by the cells from the skin, as *L. major* and *L. braziliensis*; reveal that amastigote from *L. donovani* remain naked without the protection of the PV, and internalize reach the blood stream and infect internal organs as the spleeen, liver and bone marrow, develop the visceral leishmaniasis (Tab. 2).

### Infection mechanism by amastins from *L. donovani*

The infection of host cells by *L. donovani* is based in the architecturally similarity of the amastins with Human Derlin 1 protein, which is a hydrophobic molecule compose by four transmembrane segments, similar to amastins (Fig. 9 a-b). The human Derlin 1 is a protein from the ER, that recognize and interact the human MHC-1 molecules participate in the quality control of the production of MHC-1 molecules [46], thereby is thought that amastins similar to Derlin 1, can recognize the MHC-1 molecules as host receptor from rich cells in MHC-1 molecules, such as the macrophages, dendritic cells, and Langerhans cells. The N-myristoylation sites from amastins can interact the MHC-1 molecules participate in the infection of host cells [32]. In the infection mechanism by *L. donovani* can participate the single microbody amastin slightly attach the parasite to the cell membrane of the host cells. The internalization and transport of the parasite in the infected cell is facilitated by the MHC-1 recycle and endosome formation of the infected cell, drag and transport the amastigote in the cytoplasm of the infected cell (Tab. 1).

**Fig, 9.**
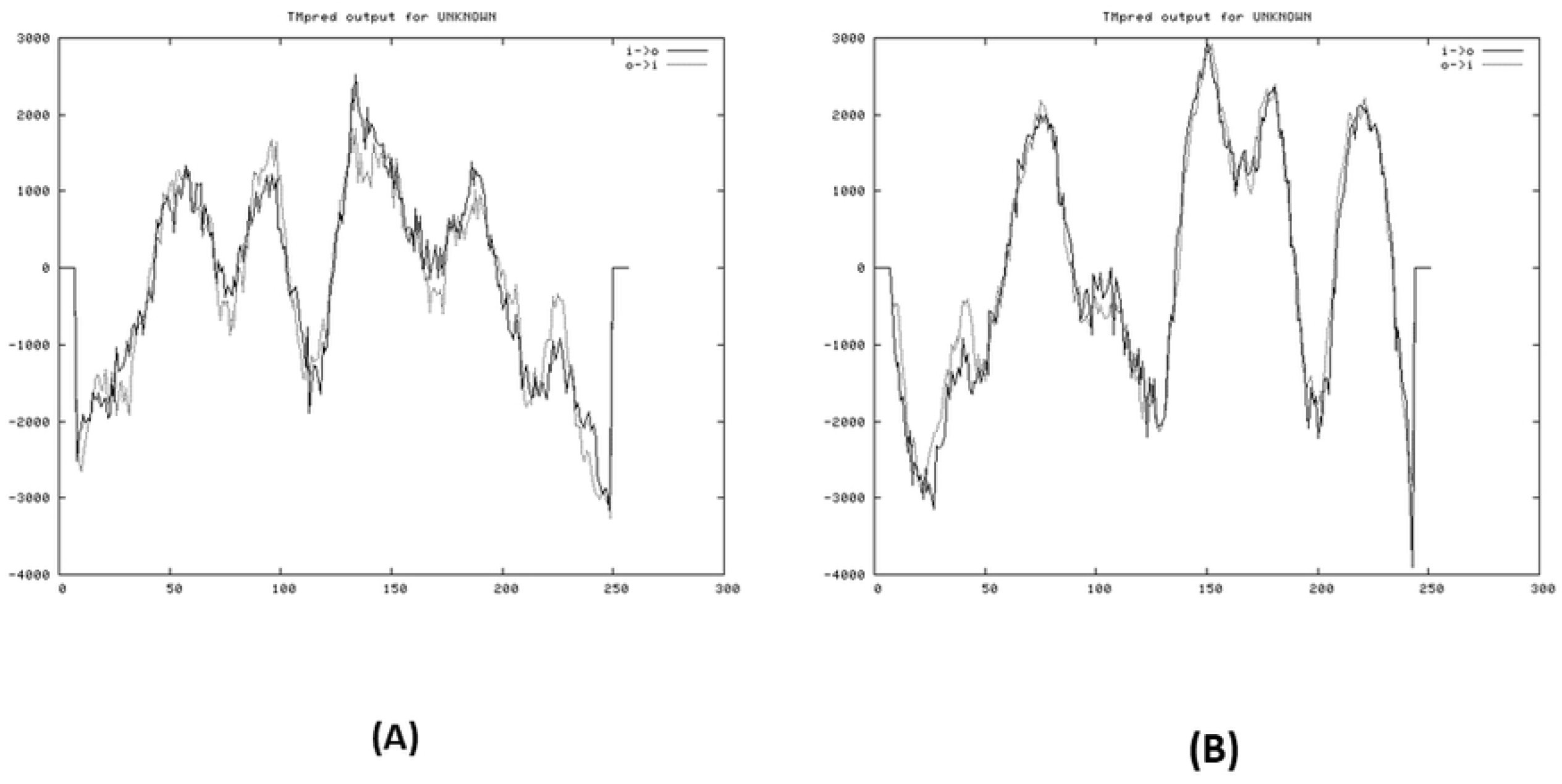
Hydrophobic patterns show the architectural similarity of these proteins. A) Human derlin1 protein B) amastin from *L. major*. Show that both proteins are hydrophobic molecules and compose by four hydrophobic regions that shape four transmembrane segments, of around 200 amino acids.

### The acid resistant mechanism of amastins from *L. donovani*

The determination of isoelectrical point (Ip) to the family of amastins from *L. donovani*, show that 20 amastins are acidic, and 30 amastins are basic. Strongly suggest that amastins from *L. donovani* developed acid resistant mechanism, that neutralize the acidic pH of the lysosomes of the infected macrophage, contribute with the deactivation of the infected macrophages, and protection of the cell membrane and membrane proteins of the parasite, allowing the develop of the amastigote, and progress of the visceral leishmaniasis (Tab. 1).

### The polarization of the immune response from protective Th1 to non protective Th2 by amastins from *L. donovani*

These results show an alternative mechanism of polarization of the immune response from protective Th1 to non protective Th2; see polarization of the immune response by *L. major* (Tab. 1).

### N-myristoylation sites from amastins from *L. donovani*

The amastins from *L. donovani* with different number of N-myristoylation sites, vary from single to 14 myristoylation sites by amastin, developed a total of 206 N-myristoylation active sites. Actually is well know protein N-myristoylation is an important fatty acylation catalyzed by N-myristoyltransferase, which is ubiquotous enzyme in eukaryotes. Specifically, atttachment of a myristoyl groups vital for proteins participating in various biological functions, including signal transduction, cellular localization, and oncogenesis. Recent studies have revealed unexpected mechanism indicating that protein N-myristoylation is invovlved in host defense against microbial and viral infection [31]; strongly indicate that N-myristoylation sites from amastins can protect the amastigote from microbial and viral infections. Actually is well know that myristoylated proteins can bind MHC-1 molecules [32]. Identification and structure of an MHC class I-Encoded protein with the potential to present N-myristoylated 4-mer peptides to T cells. [33]. Strongly indicate that N-myristoylated amastins can participate in the interaction of the human MHC-1 molecules as host receptor (Tab. 2).

### Parasite nutrition mechanism from *L. donovani*

The nutrition mechanism of the amastigote in the novel environmental conditions of the infected human cells remain unknown. These results show that diversity of the four transmembrane segments amastins can mimics carriers and transporters of four transmembrane segments [34] achieve a nutrition mechanism. The amastins derlin 1 can dissimilate the MHC-1 molecules and other subsets of molecules [35], the dissimilation products can be translocated through of the parasite cell membrane by amastins and employed as nutrient source, achieve a nutrition mechanism of the amastigote (Tab. 1).

### The family of amastins from *L. infantum*

The family of amastins from *L. infantum* is compose by 40 mature amastins (Tab. 1). The family of amastins ordered by Ip show that developed 23 molecules acididc, and 17 molecules basic. The molecular weight of the acidic Ip amastins vary from 4.44 to 6.7. The Ip of the basic amastins vary from 7.45 to 9.38. The molecular weight (MW) of the acidic amastins vary from 19.24 KD to 24.46 KD with the exception of single large acidic amastins of 57.82 KD. The MW of the basic amastins vary from 19.18 KD to 23.34 KD, with the exception of four large amastins two of 28.96 KD and 29.88 KD, and two large of 32.48 KD, and 59.14 KD (Fig. 2).

The molecular diversity of the amastins from *L. infantum* can be show in the number of transmembrane segments developed by the amastins, show that majority of the amastins of *L. infantum* developed 4 transmembrane segments (21 amastins), 11 amastins developed 3 transmemebrane segments, 5 amastins developed 6 transmemebrane segments, single amastins developed 2 transmembrane segments, and single large amastins developed 7 transmembrane segments. Show diversity of the amastins from *L. infantum* developed mainly 4 transmembrane segments (Fig. 2).

### The cellullar target of the amastins from *L. infantum*

The molecular diversity of the amastins from *L. infantum* can be see in the cellular target of the amastins, show that 28 amastins are targeted in the parasite cell membrane, 8 amastins developed the signal peptide to enter to the secretory pathway, and the mature amastins are targeted in the parasite cell membrane by 3 or 4 transmembrane segments; the results furthermore show that repertoire developed 4 non secretory proteins (Fig. 2).

### The amidation sites of the amastins from *L. infantum*

The analyses of amidation sites of the amastins from *L. infantum*, show that developed 7 amastins with 7 amidations sites, this is the intracellular trypanosomatid with amastins with more amidation sites. (See amidationa sites of the amastins from *L. donovani*) (Tab. 2).

### The N-glycosylation of the amastins from *L. infantum*

The results show that family of amastins from *L. infantum* developed 21 amastins with ASN-Glycosylation N-glycosylation site, with 31 active sites. Interestingly the results show that family of amastins did not developed none of N-linked (GlcNAc…) asparagine sites. Reveal that just one half of the amastins are ASN N-glycosylated. (See *L. donovani*.) (Tab. 2)

### cAMP and cGMP-dependent protein kinase phosphorylation sites from *L. infantum*

The repertoire of amastins from *L. infantum* developed 11 amastins with 11 cAMP and cGMP-dependent protein kinase phosphorylation sites (Tab. 2) (See *L. donovani*).

### Casein kinase II phosphorylation sites of the amastins from *L. infantum*

The results show that *L. infantum* developed 37 amastins with 94 Casein kinase II phosphorylation sites (Tab. 2) (See *L, donovani*).

### Protein kinase C phosphorylation sites of the amastins from *L. infantum*

The results show that amastins from *L. infantum* developed 35 amastins with 89 protein kinase C phosphorylation sites, vary from 1 to 8 per amastins (Tab. 2). (See *L. donovani*).

### Phosphothreonine and phosphoserine of amastins from *L. infantum*

These results show that *L. infantum* develop 1 amastin with 1 phosphoserine, and 7 phosphothreonine (**>LinJ34_V3.2660)**; whereas *L. donovani* develop numerous amastins with 137 phophoserine, and 128 phosphothreonine. Reveal that *L. infantum* is scarce phosphorylable in serine and threonine. This mechanism widely used by the amastins from *L. donovani* can be atractive for many phophorylases in serine and threonine from the infected cell contribute with the deactivation of infected cell (Tab. 2). (See *L. donovani*)

### Cell attachment sequences of amastins from *L. infantum*

The results show that family of amastins from *L. infantum* developed two amastins with the cell attachment domain with the minimal sequence of recognition RGD; and six amastins with cell attachment domain with the minimal sequence of recognition LDV. The sequence RGD is typical of integrins and the extracellular matrix. The sequence LDV is typical from the integrins from leukocytes. Strongly indicate that RGD amastins can recognize the extracellular matrix of the infected tissue such as the spleen, liver and bone marrow of the visceral leishmaniasis [36]. The amastins with the minimal sequence of recognition LDV, from leukocytes can participate in the attachment of the parasite to the macrophages, dendritic cells, and Langerhans cells, participate in the infection of the parasite. Also can interfere with the correct cellular synapsis.of activation of leukocytes such as the macrophages, and T lymphocythes [91] (Tab. 2). (See *L. donovani)*

### Microbody C-terminal cell targeting of the amastins from *L. infantum*

The results show that family of amastins from *L. infantum* developed single microbody amastins. The microbody domain is a hydrophobic C-terminal domain that recognize and passs the cell membranes [37-38]. Thereby the microbody amastins can interact and pass the cell membrane from the host cells. The single microbody amastin can participate in slightly attachment of the parasite to the host cell, contribute with the infection mechanism of *L. infantum* (Tab. 1). (See *L. donovani*).

### The N-myristoylation sites of the amastins from *L. infantum*

The results show that *L. infantum* developed 39 amastins with 129 N-myristoylation sites vary from 1 to 14 sites by molecule (Tab. 2), (See *L. donovani*.

### Amastins with aldehyde deshydrogenase active site from *L. infantum*

The family of amastins from *L. infantum* evolve two amastins with aldehyde deshydrogenase active site. The amastins with aldehyde deshydrogenanase active site can bind and mimics the human molecular environment. Actually is well know that human ALDH can unspecifically interact with biological molecules as androgen [39-40], cholesterol [41-42] and thyroid hormone [43]; as well chemical drugs as acetominophen compounds [44-45] remain unknown the biological relevance of this binding property.

These results show that membrane amastins ALDH developed by *L. infantum*, can have addditional biological functions in the human infection, can bind unspecifically host molecules as different hormones, cholesterol, and employ these molecules for mimic the human molecular environment, mask the parasite and evade the immunne response (Tab.2).

### The infection mechanism with amastins from *L. infantum*

The infection mechanism from *L. infantum* is similar to the infection mechanism developed by *L. donovani*. The infection is based in the architectural similarity of amastins with the human derlin 1 protein, which is hydrophobic molecule compose by four transmembrane segments, that recognize and interact the human MHC-1 molecules [46] (Fig. 9). Thereby is tought that amastins similar to derlin 1 protein recognize the MHC-1 molecules as host receptor, from rich cells in MHC-1 molecules, such as the macrophages, dendritic cells, and Langerhans cells. The N-myristoilation sites can bind the human MHC-1 molecules, explain the binding of the amastins to MHC-1 molecules, or cooperate with infection of rich cells in MHC-1 molecules. *L. infantum* developed single microbody amastin can participate in the attachment of the parasite to the host cell, and did not shape parasitophorous vacuole. The cell attachment with the minimal recognition site LDV can participate in the attachmen of the parasite to the host cells. These results show that are multiple ways of infection of host cells by *Leishmania*. The internalization and transport of the parasite is facilitated by the MHC-1 molecules recycle and endosome formation, drag and transport the amastigote in the cytoplasm of the infected cell.

### Polarization of the immune response from Protective TH1 to non prtective TH2 by amastins from *L. infantum*

The evasion of the immune response by polarization of the immune response from protective TH1 to non protective TH2 is similar to polarization from *L. major*. (See *L. major*).

### Parasite nutrition mechanism by amastins from *L. infantum*

The nutrition mechanism from *L. infantum* is similar to *L. donovani* nutrition mechanism. (See *L. donovani)*.

### The family of amastins from *L. major*

The *L. major* agent cause of localized cutaneous leishmaniasis (LCL) from old world. Develop the family of amastins compose by 54 mature amastins, several of them in multicopy gene (contain 37 isoforms of amastins) (Tab. 1). The table show that amastins ordered by Ip, show 27 acidic amastins and 27 basic amastins, there is no acid resistant mechanism mediated by amastins. The molecular weight of the amastins show that acidic molecules vary from MW=14.34 KD to MW=23.65 KD, with single large amastins of MW=57.56 KD. The basic amastins vary from MW=15.01 KD to MW= 22.08 KD (Fig. 3).

The cellular target of the amastins from *L. major* show diversity, reveal that 45 amastins are targeted in the cell membrane of the parasite; 2 small secretory membrane amastins of small MW= 14.34 KD and 14.39 KD they are acidic amastins. The 5 basic small secretory membrane amastins vary MW= 15.01 KD to 17.67 KD. There are two non secretory proteins of MW=23.65 KD and MW=57.56 KD (Fig. 3).

The molecular diversity of the amastins from *L. major* can be see in the number of transmembrane segments developed by these molecules, 41 amastins developed 4 transmembrane segments, 7 amastins developed three transmembrane segments; 3 amastins developed five transmembrane segments, and single amastin developed two transmembrane segments (Fig. 3).

### Amidation sites of the amastins from *L. major*

The analyses of the amidation sites of the amastins from *L. major* show that this parasite develop one amastins with one amidation site (Tab. 2). (See *L. donovani*)

### N-glycosylation amastins from *L. major*

The results show 23 amastins developed 58 ASN-Glycosylation N-glycosylation sites, reveal that *L. major* just developed 23 amastins N-glycosylated, and the majority of amastins from this parasite are not glycoproteins, these molecules can evade the immune response, antigen variation (Tab. 2) (see *L. donovani*).

### cAMP- and cGMP-dependent protein kinase phosphorylation sites of amastins from *L. major*

The results show that 5 amastins from *L. major* developed 5 cAMP- and cGMP-dependent protein kinase phosphorylation sites. (Tab. 2) (See *L. donovani*).

### Casein kinase II phosphorylation sites of the amastins from *L. major*

The results show that *L. major* contain 6 amastins developed 28 Casein kinase II phosphorylation sites (Tab, 2) (See *L. donovani*).

### Protein kinase C phosphorylation sites of the amastins from *L. major*

The results show that *L. major* contain 13 amastins developed 27 Protein kinase C phosphorylation sites. (Tab. 2) (See *L. donovani*).

### Cell attachment sequence of the amastins from *L. major*

The analyses of the cell attachment sequence in the family of amastins from *L. major* show that developed 3 cell attachment sequences with the minimal recognition sequence LDV; two are in the large amastins, and one in membrane amastin.(Tab. 2) (See *L*.*donovani*)

### N-myritoylation sites of the amastins from *L. major*

The results show that family of amastins from *L. major* develop 54 amastins with 180 N-myristoylation sites. Reveal that N-myristoylated proteins can bind the MHC-1 molecules (Tab. 2). (See *L. donovani*).

### Microbody amastins from *L. major*

Infection of host cells by *Leishmanias* becomes highly potent due to the effect of microbody amastins. Infection of the macrophages, dendritic cells, and langerhans cells by *L. major*, can be substantially improved by the expression of the 9 identical amastins called microbody amastins, which develop microbody domains in the C-terminal, the amino acids GRA, which is able of interact and pass the cell membranes [38]. The microbody amastins strongly attach the amastigotes of *L. major* to the cell membrane of the host cells (Tab. 2). The attachment with the numerous microbody amastins appears like velcro tape potency the infection (Fig. 10).

**Fig. 10.**
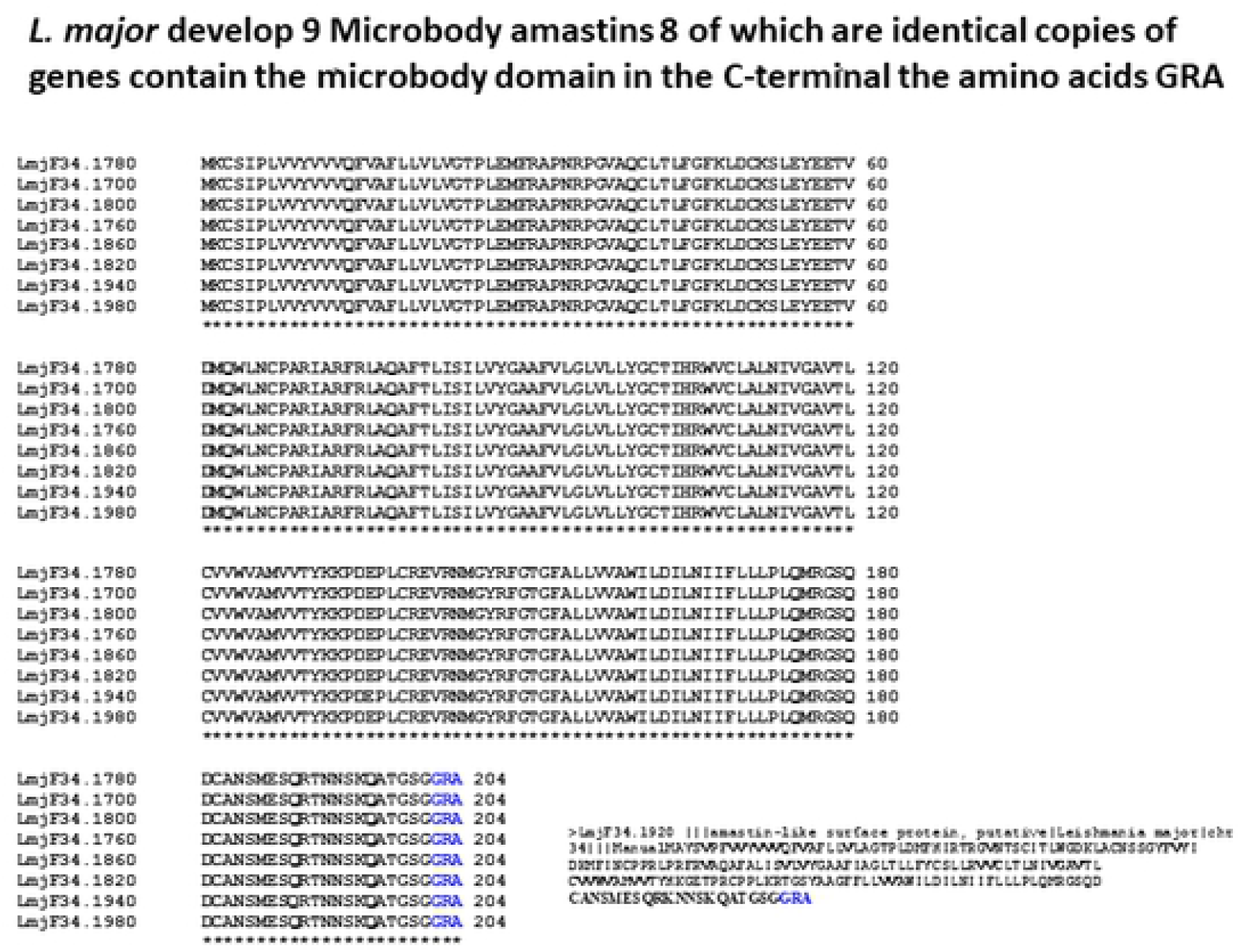
microbody amastins from *L. major* show the microbody domain in the C-terminal, the amino acids GRA. which is able of interact and pass the cell membranes. The amino acid sequence alignement was performed with eight identical copies of the amastin. The nine microbody amastin show in the lower right corner,

### Amastin laminin G domains from *L. majo*r

Laminin a component of the extracellular matrix that is found in all basal laminae and has binding sites for cell surface receptors, collagen, and heparan sulfate proteoglycan. These results show that family of amastins from *L. major* develop a membrane amastin with laminin G domain, which can interact with the collagen and heparan sulfate proteoglycan sites from the extracellular matrix of the skin tisssue, slightly attach the amastigote to the extracellular matrix (Tab. 2).

### Amastins Tyrosine sulfation site from *L. major*

Tyrosine sulfation is a post-translational modification of secreted and membrane proteins, in which a sulfate group is transferred from 3’ phosphoadenosine 5’phosphosulfate to the hydroxyl group of a tyrosine residue to form a tyrosine O4 sulfate ester. Some examples of mammalian proteins known to be tyrosine sulfataed are: complement protein C4; numerous blood coagulation proteins such as factor V. Hormones such as human choriogonadotropin α-chain; and cell surface adhesion molecules P-selectin glycoprotein ligand-1. In addition many GPCRs are tyrosine sulfated including C3a and C5a-anaphylatoxin chemotactic receptor and type 1 sphingosine 1-phosphate receptor (involved in complement activation, chemotaxis and T cell activation; follicle-stimulating hormone receptor and luteinizing hormone receptor (involved in reproductive functions, and thyroid-stimulating hormone (essential for proper endocrine function of the thyroid gland), as well several chemokines receptors, such as CCR2, CCR5, CCR8, CXCR4, CX1CR1, and DARC [47]. The family of amastins from *L. major* develop 14 amastins with tyrosine sulfation site, can mimics the complement protein C4, proteins of coagulationsuch as favtor V; cell surface adhesion molecules P-selectin glycoprotein ligand 1. Many GPCRs, including C3 and C5a anaphylotoxin chemotactic receptor and type 1 sphingosine 1-phosphate receptor (involved in complement activation, chemotaxis and T cell activation), the tyrosine sulfation site from chemokine receptor, interfere the binding of chemokines and avoid the activation of chemokine receptor, that activate the leukocytes such as the macrophages and T lymphocytes, that eliminate the parasite (Tab. 2).

### Infection mechanism by amastins from *L. major*

These results show that infection mechanism from *L. major* is based in the architectural similarity of amastins with the human Derlin 1 protein, which is a hydrophobic molecule compose by four transmebrane segments, that recognize and interact the human MHC-1 molecules [46], thereby is thought that amastins recognize the MHC-1 molecules as host recepetor for infection, from rich cell in MHC-1 molecules such as the macrophages, dendritic cells, and Langerhans cells (Tab. 1).

In the infection of *L. major* of human macrophages furthermore participate 9 microbody amastins that strongly attach the parasite to the cell membrane of the host cells, potency the infection. The N-myristoylations sites participate bind the MHC-1 molecules attach the parasite to the host cell (Tab. 2). The internalization and transport of the parasite is facilitated by the MHC-1 molecules recycle and endosome formation that drag and transport the parasite in the cytoplasm of the infected cell.

### Tropism of the amastins from *L. major* by the cells from the human skin

These results show that tropism of the amastigotes by the cells from the skin is facilitated by two molecular interactions. The first molecular interaction is facilitated by amastins that recognize and interact the human MHC-1 molecules, from rich cells in MHC-1 molecules such as the macrophages, dendritic cells, and Langerhans cells from the skin. The second molecular interaction is facilitated by microbody amastins, these numerous hydrophobic domains strongly attach the amastigote to the cell membrane of the macrophages, dendritic cells, and Langerhans cells, retain the parasite in the cells from the skin, where develop the pathogenesis of the localized cutaneous leishmaniasis (Tab. 1).

### Biogenesis of the small parasitophorous vacuoles by amastins from *L. major*

The PVs observed in some *Leishmania* are constructed by cell membrane and proteins from the endoplasmic reticulum, revealing that PV contains cell membrane from the infected cell. This implied that molecules that are trafficked to the endoplasmic reticulum are transferred to the PV and that PVs are hybrids compose of both host endoplasmic reticulum and endocytic pathway componenets [48]. *L. major* develops small and individual PVs which can fuse with the large PVs from *L. amazonensis* [49]. This work shows that the biogenesis of PV is mediated by the microbody amastins which strongly attach the parasite to the cell membrane of the host cell, shape the PV simulataneously to the infection of the host cell, and develop small and individual PVs that harbor single amastigote of *L. major* (Fig. 10).

### Novel mechanism of polarization of the immune response from protective TH1 to non protective TH2 by amastins from *L. major*

Leishmaniasis was one of the first parasitic disease which the polarization of the immune response was demonstrated, which is characterized by a shift from T helper (TH1) protective immune response to the Th2 non protective immune response [50].

Polarizationof the immune response has been successfully asssociated with several viral, bacterial, and paraste infection, and autoimmune disease in which the cytokine profile produced by one type of TH response dominates upon the other response [51-53]. In leishmaniasis, it is generally accepted that the sueptibility to *L. major* infection correlates with the dominance of the interleukin-4 (IL-4) driven by TH2 response that causes the disease; whereas the production of IL-12 and interferon-γ dominates the TH1 response that promote healing and parasite clearanace. The results of the analyses of the amastins from *L. major, L. infantum, L. braziliensis, L. donovani* and *T. cruzi* proposes a new and alternative mechanism to drive and polarize the immune response from protective TH1 to non-protective TH2 in these human parasitic diseases. This polarization can be performed by amastins with proteolytic activity, similar to the human protein derlin-1, which are essentail ER protein specilized in the degradation of MHC-1 molecules and other subsets of proteins, thereby these proteolytic amastins could degradate MHC-1 molecules and other subsets of proteins from the infected host cell [46]. The interaction of the proteolytic amastins with the MHC-1 molecules could ocurr in the PV, since it is well known that PV contain MHC-1 molecules [54] and contain ER from the host cells [55]. Loss the MHC-1 molecules by parasite degradation can results in a poor presentation of the antigenic peptides by the infected APC and inhibit the stimulation of the immune response and biological activity of the CD8^+^ cytotoxic T lymphocytes (CTL), which are the responsible for cytotoxic activity and the secretion of TH1 cytokines that stimulate the cellular response (TH1). Hence the conditions that favor the stimulation of TH2 non-protective immune response in the infected host are mantained and the synthesis patterns of their cytokines, promoted by the antigens presentation by te MHC-2 molecules in the surface of the infected cells [52-53] allow the internalization and growth of the parasite and assist the development of the parasisitic disease (Tab 1).

### Parasite nutrition by amastins from *L. major*

The molecular diversity of the numerous four transmembrane segments amastins can mimics the four transmembrane segments transporters and carriers achieve an nutrition mechanisms of the parasite cell membrane. The amastins can dissimilate the human MHC-1 molecules and the disssimilation products can be translocated through of the parasite cell membrane and employed as nutrition source develop an nutrition mechanism [46] (Tab 1). (See L. *donovani*)

### The family of amastins from L*eishmania braziliensis*

The family of amastins from *L. braziliensisis* is compose by 47 mature amastins; the sequence >LmjF31.0450; the sequence >LbrM18_V2.0450 |phosphatidic acid phosphatase; the sequence >LbrM08_V2.1040 |beta tubulin; and the sequence >LbrM20_V2.1090 |kinesin were discarded from the family of amastins from *L. braziliensis*. The sequence >LbrM08_V2.1100 without name was taked as amastin by the large similarity with the sequences of the family of amastins from *L. braziliensis* (data not shown).

The amastins from *L. braziliensis* ordered by Ip show that 21 amastins are acidic, and 26 are basic molecules; there is no clear acid resistant mechanism.

The MW of acidic amastins vary from 20.95 KD, to 23.9 KD; The MW of basic amastins vary from 14.93 KD, to 26.5 KD with the exception of 3 large amastins of 34.19 KD, 49.72 KD, and 58. 79 KD; 2 are non secretoery proteins (intracellulars), and one developed the signal peptide, pass by the secretory pathway and the mature protein is targeted in the parasite cell membrane, by five transmembrane segments (Fig. 4).

### The transmembrane segments of the amastins from *L. braziliensis* show different distribution

The molecular diversity of the amastins from *L. braziliensis* can be appreciated in the number of transmembrane segments developed by the amastins, is generally accepted that amastins are hydrophobic molecules composed by four transmembrane segments. These results show that 21 amastins developed 4 transmembrane segments; 24 amastins developed 3 transmembrane segments; 2 amastins developed 5 transmembrane segments, and small amastins of 135 amino acids developed two transmembrane segments, this molecule can be the evolutionary traces of the creation of the diversity of the repertoire of amastins from *L. braziliensis*. Conclude that majority of amastins from *L. braziliensis* developed three transmemebrane segments, the biological relevance of the amastins of three transmembrane segments amastins from *L. braziliensis* remain unknown, but the overexpresssion of amastins can shape protective protein cover in the parasite (Fig. 4).

### The amidation sites of the amastins from *L. braziliensis*

The analyses of the amidation sites of the amastins from *L. braziliensis* show tha this parasite develop two amastins with two amidation sites. (See *L. donovani*).

### ASN-Glycosylation N-glycosylation site of the amastins from *L. braziliensis*

The results show that *L. braziliensis d*eveloped 25 amastins with 47 N glycosylation active sites. Reveal that half of amastins from *L. braziliensis* are glycosylated (Tab. 2) (See *L. donovani*).

### cAMP- and cGMP-dependent protein kinase Phosphorylation sites of the amastins from *L. braziliensis*

The results show that *L. braziliensis* develope 13 amastins with 18 cAMP- and cGMP-dependent protein kinase phosphorylation sites. (Tab. 2) (See *L. donovani*)

### Casein kinase II phosphorylation sites of the amastins from *L. braziliensis*

The results show that 40 amastins developed 114 Casein kinase II phosphorylation sites (Tab. 2). (See *L. donovani*)

### Protein kinase C phosphorylation sites of the amastin from *L. braziliensis*

The results show that *L. braziliensis* developed 38 amastins with 78 Protein kinase C phosphorylation sites (Tab. 2). (See *L. donovani*).

### N-myristoylation sites of the amastins from *L. braziliensis*

The results show that *L. braziliensis* develop 42 amastins with 157 N-myristoylation sites, strongly indicate that N-myrystoilation sites play important role in the binding of MHC-1 molecules (Tab. 2) (See *L. donovani*).

### Phosphoserine and phosphotreonine of the amastins from *L. braziliensis*

Amazingly the family of amastins from *L. braziliensis* did not developed none phosphoserine or phosphothreonine, whereas *L. donovani* develop 137 phosphoserine, and 128 phosphothreonine, strongly indicate that amastins from *L. donovani* are widely phosphorilable, particicpate in the deactivation of infected cell (Tab. 2).

### Infection mechanism of the amastins from *L. braziliensis*

The infection mechanism from *L. braziliensis* is based in the architectural similarity of amastins and the human derlin 1 proteins, both proteins are hydrophobic and compose by four transmebrane segments. The Derlin1 protein recognize and interact the human MHC-1 molecules, is tought that amastins recognize the MHC-1 molecules as host receptor for infection of cells rich in MHC-1 molecules such as the macrophages, dendritic cells, and Langerhans cells. The amastins from *L. braziliensis* developed 42 amastins with 157 N-myristoylation sites which can bind the MHC-1 molecules. The *L. braziliensis* develop 3 microbody amastins that attach the parasite to the host cell potency the infection (Tab. 1). The internalization and transport of the parasite in the cytoplams of infected cell is facilitated by the MHC-1 recycle and endosome formation drag and transport the parasite in the cytoplasm of infected cell.

### The polarization of the immune response from protective TH1 to non protective TH2 by amastins from *L. braziliensis*

It is similar to the polarization of the imune response from protective TH1 to non protective TH2 from *L. major* (Tab. 1). (See *L. major)*

### The parasite nutrition mechanism of the amastins from *L. braziliensis*

The parasite nutrition mechanism from *L. braziliensis* is similar to the *L. donovani* nutrition mechanism (Tab. 1). (See *L. donovani*).

### The family of amastins from *Trypanosoma cruzi*

The family of amastins from *T. cruzi* is compose by 12 mature amastins (Tab. 1), all amastins are hydrophobic and basic molecules, their MW vary from 10.39 KD to 21.43 KD, the small isoform of 96 amino acids, show the evolutionary traces of the creation of the repertoire, all amastins are targeted in the cell membrane of the parasite, 11 amastins developed 4 transmembrane segments, with the exception of small isoform that is compose by 1 transmebrane segment (Fig. 5).

### Acid resistant mechanism by amastins from *T. cruzi*

All amastins from *T. cruzi* are basic, strongly indicate that amastigote is embebebd in acidic environment, as can be the acidic environent of the infected macrophages, the overexpresssion of the amastins neutralize the lysosomal activity deactivate the infected macrophage (Tab. 1; Fig. 5).

### The amidation sites of the amastins from *T. cruzi*

The analysese of the amidation sites of the amastins from *T. cruzi* show that did not developed amidation sites, strongly indicate that amastigotes are internalized and transported by the MHC-1 recycle and endosome formation drag and transport the parasite in the cytoplams of the infected cells (Tab. 2). (See *L. donovani*).

### The cAMP- and cGMP-dependent protein kinase phosphorylation sites of the amastins from *T. cruzi*

The family of amastins from *T. cruzi* developed 3 amastins with three cAMP- and cGMP-dependent protein kinase phosphorylation sites. (Tab. 2). (See *L. donovani)*.

### The casein kinase II phosphorylation sites of the amastins from *T. cruzi*

The family of amastins from *T. cruzi* developed 8 amastins with 21 Casein kinase II phosphorylation sites (Tab. 2), (See *L. donovani*).

### Protein kinase C phosphorylation sites of the amastins from *T. cruzi*

The family of amastins from *T. cruzi* develop 11 amastins wit 21 Protein kinase C phosphorylation sites (Tab. 2). (See *L. donovani*).

### N-myristoylation sites from amastins from *T. cruzi*

The twelve amastins developed 30 N-myristoylation sites, reveal that amastins can react with T cells, and bind the MHC-1 molecules (Tab. 2) (See *L. donovani*).

### N-glycosylation from amastins from *T. cruzi*

The twelve amastins did not developed N glycosylation sites, amazinzgly did not developed none glycosylation site, none amastin from *T. cruzi* is glycoprotein (Tab. 2). (See *L: donovani*)

### The infection mechanism by amastins from *T. cruzi*

The infection mechanism from *T. cruzi* is based in the architectural similairity of amastins and the human derlin-1 protein, which is a hydrophobic moleculle compose by four transmembrane segments, that recognize and interact the human MHC-1 molecule, thereby is touched that amastins can recognize the human MHC-1 molecules as host receptor from rich cells in MHC-1 molecules, such as the macrophages, dendritic cells, and langerhans cells. The twelve amastins developed 30 N-myristoylation sites, which can bind the MHC-1 molecules (Tab. 1, 2). The internalization and transport of the parasite is facilitated by the MHC-1 molecules recycle and endosome formation that drag and transport the parasite in the cytoplams of the infected cell.

### The polarization of the immune response from protective TH1 to non protective TH2 from *T. cruzi*

It is similarto to polarizationa of the immune respose from *L. major* (Tab. 1) (See *L. major*).

### The parasite nutrition mechanism of the amastins from *T. cruzi*

It is similar to the mechanism of nutrition from *L. donovani* (Tab.1). (See *L. donovani*).

## Conclusions

The large families of amastins developed by *L. donovani, L. infantum, L. major, L. braziliensis*. and *T. cruzi* are strongly associated with the develop of intracellular parasitism from rich cells in human MHC-1 molecules such as the human macrophages, dendritic cells, and Langerhans cells

The amastins are similar to the human derlin 1, both proteins are hydrophobic molecules compose by four transmembrane segments, of around 200 amino acids.

The amastins recognize and interact the human MHC-1 molecules as main host receptor from rich cells in MHC-1 molecules such as the macrophages, dendritic cells, and Langerhans cells.

The microbody amastins potency the infection of host cells as co receptors.

The tropism of *L. major* and *L. braziliensis* by the cells from the skin is facilitated by two molecular interactions, the first molecular interaction is facilitated by te amastins iteract the human MHC-1 molecules, and the second molecular interaction is facilitated by the microbody amastins that interact and pass the cell membrane, retain the parasite in the cells from the skin. *L. donovani* and *L. infantum* developed single microbody amastins, and *T. cruzi* did not developed none microbody amastins, thereby cannot be retained by the cells from the skin pass to the blood stream and infect internal organs.

The microbody amastins participate in the biogenesis of the small PV from *L. major* (9 microbody amastins numeorus molecular interactions), and the large PV from *L. braziliensis* (3 microbody amastins few molecular interactions).

Acid resistant mechanism by basic amastins developed by *L. donovani* and *T. cruzi*.

All the human Trypanosomatids intracellulars develop internalization and transport of the parasite by the MHC-1 molecules recycle and endosome formation drag and transport the parasite in the cytoplasm of infected cell

All the human intracellular Trypanosomatids developed evasion of the immune response by polarization of the immune response from protective TH1 to non protective TH2.

All the human intracellular Trypanosomatids develop the nutrition mechanism by dissimilation of MHC-1 molecules and translocation of the dissimilation products trough of the parasite cell membrane, and employed as nutrient source.

All amastins developed few amidation sites include *T. cruzi* did not developed none amidation site, did not play relevant biological function in these parasites (Tab. 2).

All the parasite intracellular except *T*.*cruzi* develop amastins with ASN-glycosylation N-glycosylation site *L. donovani* develop 38 amastins with 38 active sites, The rest of parasites veloped one half of glycocylated amastins with mutiple active sites. Just *L. donovani* developed 45 amastins with 45 N-linked (N acetil glucosamida) asparagine, show that *L*.*donovani* is the parasite with more N-glycosylated amastins; perform more antigen variation, and evasion of the immune response

All the parasites developed amastins with cAMP and cGMP dependent protein kinase phosphorylation site, however *L. braziliensis* develop more amastins (13) with 18 active sites

All amastins from these intracellular parasites develop the Casein kinase II phosphorylation sites, *L. braziliensis* develop 40 amastins with 114 active sites.

All amastins from these parasites developed Protein kinase C phosphorylation site, however *L. infantum* developed the majority of amastins develop 35 amastins with 89 active sites and *T. cruzi* 11 amastins with 22 active sites, being the parasites with more amastins and active sites.

Just *L. donovani* developed 2 amastins with 2 Tyrosine kinase phosphorylation site 1.

All parasites developed phophoserine phosphothreonine exept *L. braziliensis, L donovani* develop 137 phosphoserine and 128 phosphothreonine, followed by *L. major* tha develop 14 and 4 respectively. Reveal that all these parasites can deactivate in different grade the infected cell, interfere the phosphorylation signals of activation of the infected cell.

All these parasite intracellular developed numerous amastins with N-myristoylation sites, *L. donovani* is the parasite with more N-myrystoylation sites develop 48 amastins with 206 active sites, reveal that all these parasites can widely interact the MHC-1 moleules.

The cell attachment sequence with the minmal sequence of recgnition RGD and LDV were developed by *L. donovani, L. infantum* and *L. major*, strongly indicate that can particicpate in the infection mechanism.

The Aldehyde deshydrogenase cysteine active site was developed by two amastins from *L: infantum*, strongly indicate thst can molecular mimmics, components of the infected cell.

Laminin G domain was developed by single amastins from *L. major*, indicate that amastins from *L. major* can attach the parasite to the extracellular matrix.

Tyrosine sulfation sites developed by 14 ammastins from *L. major*, indicate that can interfere the binding of the chemokines, avoid te activation of the immune response by chemokines.

### Evolutionary implications

#### The evolution theories

The genial theory of evolution of species coined by Charles Darwin and Alfred Wallace in 1858 [56] Darwin 1859 [57] was based in the analysese of morphological similarities of the development of different embryos to adult from diverse living species, and comparative analyses of fossils. They postulated that diversity of the species that inhabit the earth, oceans and fresh water evolved from worms to man. The human been descend from primates as chimpanzee and gorilla; and all organisms of the earth growth over the effects of the natural selection adapted their specific environments.

The modern theory of evolution postulates that changes of the evolution of the organism are fixed in the DNA, and the accumulation of the small random variation (mutations), recombination of DNA, and the ordering of this genetic variation by natural selection, is the cause of evolution of the species; as well the macroevolutionary process and speciation can be explained by genetic mechanisms [58].

The determination of age of the marine microfossil indicate, that life was originated in the sea approximately 388 mya, developing all the diversity of microscopic and macroscopic organisms inhabit the oceans [59] which eventually invade and colonize the earth, shape the diversity of the species, from microscopic organism such as bacteria prototozoan, algae, invertebrates to superior greeen plants and mammals [60-61]. Actually there is large body of morphological and molecular evidences that support the evolution of species alomg of the phylogenetic scale. The evolution of species additionally is supported by numeorus pieces of fossils, identify many related or close species, as well as many extinct species, that indicate the origin of new species by evolution and their loss by natural selection [62] in similar way fossils with pre-hominid features have been found, with the help of computer models, have been possibly produce the human phylogenteic tree that show the macroscopic morphological features of the evolution of the man.

There are additional theories regard the origin of DNA and species, such as the selfish gene, which talk that genes were slowly created by the cells, and distributed and preserved in the species by millions of years. The equilibrium punctuated talk that contrary of slow evolution, talk that evolution should be rapid. The junk DNA talk that genomes contain many sequences that did not achieve biological fucntion called junk DNA.

The bacteria are the living organisms more employed for invetsigations about molecular evolution, is clear that bacteria evolved by changes in the DNA sequences, as micromutations, recombination of genes, gene duplication, and gene transfer from bacteria to bacteria, and gene transfer from host to bacteria [63], contributing with the creationof new strains and evolution of new species [64].

Spite of the numerous investigations of molecular evolution of bacteria, and more complex phenotypic species, there are not reports that documented at the level of DNA, the molecular changes that promoted the origin and evolution of the novel genes, novel proteins multiple cells and organisms.

The evolution and develop of intracellular parasitism by amastins from *L. major, L. infantum, L. braziliensis, L. donovani*, and *T. cruzi* performed in this article can be employed as excellent model for the study of invasion, evolution, adaptation, and speciation of the organisms to new environment.

Actually have beeen postulated that parasitism of the numerous species of parasites arise by single mutation of the gene, accumulation of micromutations, or gene transfer from the host to parasite, mimics molecular features of the host, that facilitate the installation and develop of the parasitism [65-67]. In similar way actually it is well know that parasites have developed diverse mechanism for evasion of the immune response [68-69].

These results show the evolution of intracellular parasitism from *L. donovani, L. infantum, L. major, L. braziliensis*, and *T. cruzi* by the families of amastins, were created by genomic response.

### The complexity of the genomic response from human intracellular Trypanosomatids can be synthesized in 17 main features (1-17)

1. The genomic response ocurr in multiple genes from a family of genes or multiple families of genes, from single chromosome or multiple chromosomes of the genome, with the end of generate molecular changes, novel genes, novel active sites, and novel biological functions of the cells that resolve the novel enviromental challenges.
2. The genomic response allows investigate the molecular changes and process that participate in the creation of novel genes, and develops of novel biological function of the genes, and genomes of the cells, as well the origin and evolution of atypical molecules of the genome.
3. The genomic response allow identify the evolutionary strategies employed to shape the molecular and functional diversity of the large families of proteins that contribute with the variety of successful biological functions.
4. The genomic response employs multiple evolutionary strategies to create diversity of proteins, to shape multigene family of proteins, or different multigene families of proteins that resolve the novel environmental challenges, allow the survival, growth, and replication of the cell.
5. The genomic response employing the bioinformatical tools, allow investigate, and to know the molecular and functional features from multiple proteins, and large families of proteins, and genomes, reveal the biological role of the large synthesis of DNA, and diversity of sequences, and novel genes, families of genes, and genomes of the cells, which are impossible of investigate by the actual experiemental and theoretical methods.
6. Based in the results of these analyses, the genomic response involve the particicpation of complexes of multiple molecules in the develop of the genomic response as DNA enzymes, DNA polymerases, DNA topoisomerases, DNA gyrases, RNA polymerases, mRNA, etc., [70-75], that participate in the synthesis of large amounts of DNA, achieve mutations, creating start codons, achieve internal expansion of DNA, copy and insert DNA, and allow the growing of DNA, endonucleases of restriction, etc. transcription of DNA, and translation of mRNA.
7. The genomic response is induced by novel environmental challenges that affect the parasite cell (novel mammal host, as novel insect vector, drugs anti-parasite); and drastic environmental challenges similar to disruption of the environment as the radioactivity (as the challenges from the immune response from human. Based in the results of this article the genomic response is induced by novel environmental challenges; that interfere with the correct mechanisms of survival, growth, and replication; which the cell response redirect the biological, metabolic and energetic forces of the cells to develop the genomic response to resolve the novel environmental challenges.
8. The data show that genomic response that creatests the large family of retrotransposons, trans-sialidases, from *T. cruzi*, occur mainly in the subtelemeric region o end of telomeric region of the chromosomes [76-77]. Thereby the telomerase from *T. cruzi* could play pivotal role develops of genomic response; and the particicipation of additional or DNA enzymes, modify the sequences to create the novel genes, and perform changes in novel genes.
9. The genomic response is induced by novel enviromental challenges that affect the survival of the cell (as novel insect vectors, anti-parasite drugs); and drastic environmental challenges similar to disruption of the environment as the radioactivity, the challenges from the immune response from human and multiple animal hosts, novel insect vector and drugs anti parasite). Based in the results of this article the genomic response is induced by novel environmental challenges that interfere with the correct mechanism of survival, growth and replication of the cells; which the cell response redirect the biological, metabolic, and energetic forces of the cells to develop the genomic response to resolve the novel environmental challenges.
10. The data show that genomic response that createst the large family of retrotransposons from *T. cruzi*, occurr mainly in the subtelomeric region or end of telomeric region of the chromososme [76-77]. Thereby the telomerase from *T. cruzi* could play pivotal role develops the genomic response and the participation of additional or DNA enzymes, modify the sequences to create the novel genes, and perform changes in the novel genes.
11. The genomic response can end when the synthesis of large amount of novel DNA, enclose the develop of successful genes, proteins and develop novel biological functions, gene arranges, and novel mechanisms of regulation of the expresion of genes, developed novel genotypes and phenotypes of the cells, novel strains, or novel species of the cell, which can follow different evolutionary routes that resolve the novel environmental challenges, allow the growth and replication of the cells or organism; in the case of these investigations after the genomic response arise follow novel differentiation of parasite appear the amastigote from these parasites, and novel parasite life cycle, and developed different and novel human parasitic disease.
12. The analyses of this article identify several and independent genomic responses, which occur depending of the number of novel and different environemental challenges, which were resolved during the natural historyof the life of each cell or organism.
13. The results of these analyses show that genomic response of the families of amastins from *L. major, L. infantum, L. braziliensis, L. donovani* and *T. cruzi* synthetize large amount of novel DNA, develop large families of amastins, which did not develop large number of immature genes and pseudogenes (Tab 1). However there are families of genes such as the mucins, retrotransposons, dispersed gene family, trans-sialidases, Gp63 from *T. cruzi* (articles in process of publish), which develop large amount of immature DNA and pseudogenes, strongly indicate that these sequences don’t have apparent biological role, which could be considered as junk DNA. The genomic response from this work, show that is necessary large synthesis of novel DNA to resolve the novel challenges, produce rapidity successful genes that resolve the novel environmental challenges. However in the genomic response can remain in the genome many immature sequences and pseudogenes, which form part of the architecture and structure of the chromosomes and nucleus of the parasite. The evolution by genomic response depend of the environmental challenges response with different amounts of DNA. Finally in the case of the dangerous envrironmental challenges of the cell, the cells response with large syntheisis of DNA from the genomic response similar as retrotransposons [78-79]. As conclusion based in this analyseses the junk DNA, is immature DNA that remain in the genomes as evolutionary traces of the induction of the genomic response, form part of the large families of succesful proteins, this way is best form of create large families of proteins to resolve environmenetal challenges.
14. Based In these analyses, in the genomic response there are genes encode for mature proteins, which could appear don’t have relevant biological role, which could appear junk DNA, however based in the results of these analyses can achieve redundant biological functions or the determination of additional analyses employ the bioinformatical tools can help to stablish biological role as Ip/pH of the cell contribute with the homeostasis of the parasite. Based in these results there is not junk DNA, since is the only way synthesize large amount of DNA, and large families of proteins by genomic response.
15. The evolution by genomic response is in accord with the theory of equilibrium punctuated [80], which talk that macroevolution and microevolution of the species should be rapid, select the most successful genotypes. The results of this article allow co probate the theory of equilibrium punctuated show 5 clear examples of human intracellular Trypanosomatids that show the the rapid genomic response, and other 4 examples of the parasite Plasmodium (articles in process of publish) that show the rapid genomic response, open the new complex and rapid evolution called here genomic response.
16. The genomic response still present species evolutionary active, there are frequent examples of asexual reproduction species, like this article, and sexual reproduction organisms, which mutant and produce new species [81-84], however is important find mutant organism of the opposite sex which cross and produce fertile offspring and novel species (Speciation).
17. The genomic response produce novel, coherent, inclusive explanation regards origin and meaning of the genes, the origin of typical and atypical molecules, origin of the genomes, origin of life, origin of eukaryotes and prokaryotes cells; virus, fungus, parasites, yeast, protozoan, metazoan, etc., and all types of cells from different environment of the planet earth. In similar way the evolution by genomic response can explain the originand evolution of superior organisms, and asexual organism from the different environment, as well as the origin of sexual reproduction and complex phenotype organism, and atypical or mutant organism. The genomic response produce novel explanation for multiple concepts as speciation, hybrids, and explain the existence of multiple phylogenetic trees of life, and the origin of life.

## Notes

### Competing Interest Statement

The authors have declared no competing interest.

